# Systems genetics analysis of human body fat distribution genes identifies Wnt signaling and mitochondrial activity in adipocytes

**DOI:** 10.1101/2023.09.06.556534

**Authors:** Jordan N Reed, Jiansheng Huang, Yong Li, Lijiang Ma, Dhanush Banka, Martin Wabitsch, Tianfang Wang, Wen Ding, Johan L.M. Björkegren, Mete Civelek

## Abstract

**BACKGROUND:** Excess fat in the abdomen is a sexually dimorphic risk factor for cardio-metabolic disease. The relative storage between abdominal and lower-body subcutaneous adipose tissue depots is approximated by the waist-to-hip ratio adjusted for body mass index (WHRadjBMI). Genome-wide association studies (GWAS) identified 346 loci near 495 genes associated with WHRadjBMI. Most of these genes have unknown roles in fat distribution, but many are expressed and putatively act in adipose tissue. We aimed to identify novel sex- and depot-specific drivers of WHRadjBMI using a systems genetics approach.

**METHODS:** We used two independent cohorts of adipose tissue gene expression with 362 - 444 males and 147 - 219 females, primarily of European ancestry. We constructed sex- and depot-specific Bayesian networks to model the gene-gene interactions from 8,492 adipose tissue genes. Key driver analysis identified genes that, in silico and putatively in vitro, regulate many others, including the 495 WHRadjBMI GWAS genes. Key driver gene function was determined by perturbing their expression in human subcutaneous pre-adipocytes using lenti-virus or siRNA.

**RESULTS:** 51 - 119 key drivers in each network were replicated in both cohorts. We used single-cell expression data to select replicated key drivers expressed in adipocyte precursors and mature adipocytes, prioritized genes which have not been previously studied in adipose tissue, and used public human and mouse data to nominate 53 novel key driver genes (10 - 21 from each network) that may regulate fat distribution by altering adipocyte function. In other cell types, 23 of these genes are found in crucial adipocyte pathways: Wnt signaling or mitochondrial function. We selected seven genes whose expression is highly correlated with WHRadjBMI to further study their effects on adipogenesis/Wnt signaling (ANAPC2, PSME3, RSPO1, TYRO3) or mitochondrial function (C1QTNF3, MIGA1, PSME3, UBR1).

Adipogenesis was inhibited in cells overexpressing ANAPC2 and RSPO1 compared to controls. RSPO1 results are consistent with a positive correlation between gene expression in the subcutaneous depot and WHRadjBMI, therefore lower relative storage in the subcutaneous depot. RSPO1 inhibited adipogenesis by increasing β-catenin activation and Wnt-related transcription, thus repressing PPARG and CEBPA. PSME3 overexpression led to more adipogenesis than controls. In differentiated adipocytes, MIGA1 and UBR1 downregulation led to mitochondrial dysfunction, with lower oxygen consumption than controls; MIGA1 knockdown also lowered UCP1 expression.

**SUMMARY:** *ANAPC2, MIGA1, PSME3, RSPO1,* and *UBR1* affect adipocyte function and may drive body fat distribution.

## Introduction

Obesity (measured as body mass index (BMI)) is a condition that affects roughly 40% of Americans^1^ and significantly increases the risk of cardio-metabolic disease^2,3^. Excess energy is primarily stored as lipid in two large adipose tissue depots, abdominal visceral and lower-body subcutaneous, where adipocytes in each depot can expand by increasing their number (through pre-adipocyte proliferation and differentiation) or their size (through adipocyte glucose and free fatty acid uptake, lipogenesis, and lipolysis) to increase storage capacity^4,5^. While subcutaneous adipocytes expand to store lipids safely and efficiently^6^, visceral adipocytes are less able to accommodate excess energy through adipocyte expansion^7,8^, which causes inflammation^9–11^, insulin resistance^12^, hypertension^13^, and negative effects on lipid metabolism^14^ in other tissues. Thus, disproportionate visceral fat storage contributes to the disease risk imparted by over-nutrition; multiple studies find that the ratio of waist circumference to hip circumference (WHR), an approximation of human fat distribution, and WHR independent of overall obesity (WHRadjBMI), are both similar or better predictors for cardiovascular disease risk and Type II Diabetes risk than BMI^15–17^. Males generally have higher WHRadjBMI and higher disease risk than females^6,18,19^. Our understanding of the genetic and molecular mechanisms that contribute to fat distribution is limited, and thus unlike overall obesity^20,21^, no targeted therapeutics or lifestyle interventions to combat abdominal obesity are known^22^.

To date, only five genes (*KLF14, LRP5, TBX15, RSPO3*, *SHOX2*) are mechanistically linked to disproportionate fat storage in one depot compared to the other. All of them affect pathways crucial to the expansion of subcutaneous and visceral adipocyte populations^23–29^. *LRP5* and *RSPO3* affect adipocyte differentiation by controlling Wnt signaling, while *TBX15* controls adipocyte differentiation and mitochondrial function.

The Wnt signaling pathway is a well-established driver of cell fate, differentiation, and proliferation in many cell types^30^; Wnt inhibits adipogenic differentiation by transcriptionally upregulating osteogenic genes while downregulating *PPARɣ* and *CEBPα*^31^. In many contexts, the non-canonical Ca^2+^ form of Wnt signaling is inhibitory of the canonical Wnt pathway^32^. Wnt signaling activity is positively associated with visceral adiposity^33,34^, and many Wnt pathway genes, especially ligands and receptors used in Ca^2+^ non-canonical Wnt signaling, are differentially expressed between fat depots^35^.

Mitochondrial function correlates strongly with cardio-metabolic diseases and can alter adipogenic differentiation^36^. In mature adipocytes, mitochondria can facilitate physical connections with the lipid droplet^37^, dissipate excess energy via thermogenesis through UCP1^38^, and promote lipid homeostasis^36^. Human visceral fat has increased mitochondrial activity compared to subcutaneous fat^39^, and in multiple metabolic disease states, only visceral mitochondria become dysregulated^39,40^. We hypothesize that other putative drivers of fat distribution affect Wnt signaling or mitochondrial function in adipocytes, with different outcomes in each depot.

Many genes potentially contribute to fat distribution. WHRadjBMI is a complex trait that is up to 60% heritable^17^, and recent genome-wide association study (GWAS) meta-analyses have uncovered ∼350 loci associated with WHRadjBMI. Approximately one third of the GWAS loci are associated with fat distribution in only one sex^41,42^, and some have greater magnitude of association in one sex^43^. Many of the ∼500 genes nearby these loci are have active regulatory regions in adipose tissue^44^ and are highly expressed in adipose tissue, many with differential expression between depots^45,46^. The expression levels of many of these genes are associated with genetic variants in within the loci, which are termed expression quantitative trait loci (eQTLs)^47–49^. Single-gene defects that cause lipodystrophy, an extreme fat distribution phenotype, also affect adipocyte function^50^.

Computational models that describe gene-gene interactions, such as networks, can be powerful tools for interrogating regulatory relationships between many genes at once and prioritizing biologically relevant regulators in the geneset^51,52^. Correlation-based networks, which identify groups of co-regulated genes^53^, are frequently used to nominate disease regulators, as well as to annotate genes of unknown function or identify conserved gene programs^54^. Bayesian networks improve upon correlation-based networks by modeling causal, directed connections, and have been used to understand gene-gene interactions in many human disease models^54–62^. The directed network structure allows researchers to easily nominate *in silico* ‘key drivers’ of tissue-specific gene regulation and of disease^67^. Network key driver genes are putatively regulators of tissue- or disease-relevant processes. While this process allows us to prioritize certain GWAS candidate genes over others, it can also nominate candidate regulators that are not genetically regulated. In various cell types and disease models, these key driver genes, both GWAS candidates and other mechanistic genes, are biologically relevant and alter disease-related pathways *in vitro*^55–66^.

To identify and prioritize candidate genes involved in fat distribution, we harnessed the predictive power of the Bayesian network construction and key driver analysis approach to identify genes likely to drive fat storage in subcutaneous and in visceral fat (Figure 1). Due to the observed differences in body fat distribution between males and females, we constructed separate Bayesian networks of each sex-depot pair to model the distinct gene regulation in each tissue. To increase the predictive power of key driver analysis, we identified key driver genes replicated in two independent cohorts and used additional publicly available data to prioritize key driver genes expressed in (pre-)adipocytes, unstudied in adipose tissue, and associated with fat storage in humans or mice. We focused on prioritized key drivers that affect Wnt signaling or mitochondrial function in other cells, and we prioritized seven genes functionally profile in subcutaneous pre-adipocytes.

**Figure 1:**
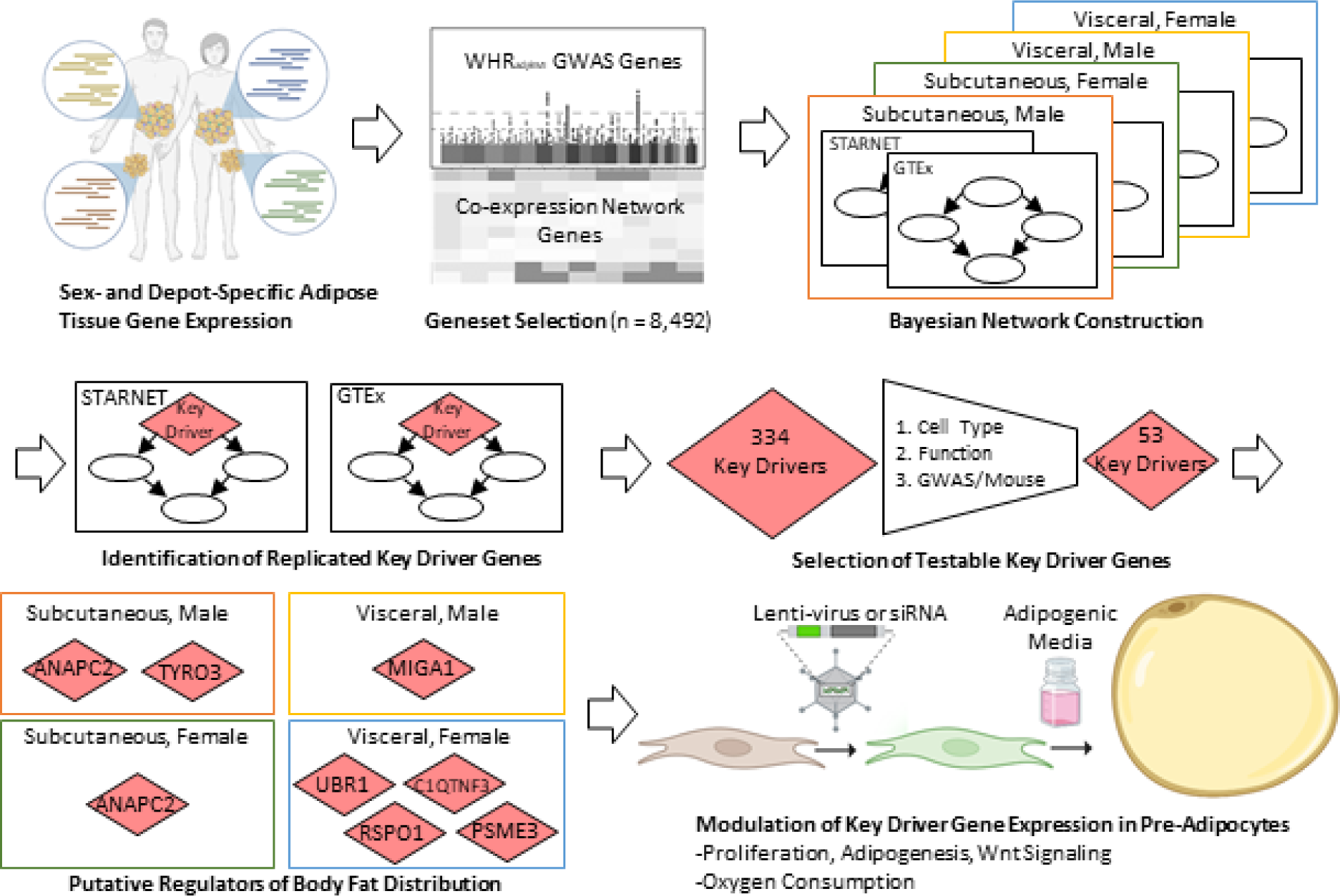
Overview of network construction, key driver gene identification, and functional validation: Publicly available RNA-seq gene expression data (GTEx^68^ and STARNET^69^) from subcutaneous and visceral adipose tissue were subsetted between males and females. Co-expression network genes and WHR_adjBMI_ GWAS^41^ genes used to construct Bayesian networks representing each sex and depot using RIMBANET^70^. Key driver genes shared between STARNET and GTEx were identified in each network. 53 key driver genes expressed in (pre-)adipocytes but unstudied in adipose tissue were prioritized for further study. Seven selected key driver genes identified were perturbed in human pre-adipocyte cells, and functional readouts of adipogenesis, Wnt signaling, proliferation, and mitochondrial oxygen consumption were collected.

## Results

### Bayesian Networks model adipose tissue gene connections in a sex- and depot-specific manner

We interrogated two independent datasets of subcutaneous and visceral adipose tissue gene expression, Genotype-Tissue Expression project (GTEx)^68^ and the Stockholm-Tartu Atherosclerosis Reverse Network Engineering Task (STARNET)^69^ (Methods) and stratified each dataset by sex. There are about twice as many males as females in each resulting dataset (Table 1).

**Table 1:** Adipose tissue donor characteristics.

|  | Subcutaneous |  |  |  | Visceral |  |  |  |
| --- | --- | --- | --- | --- | --- | --- | --- | --- |
|  | Male |  | Female |  | Male |  | Female |  |
|  | STARNET | GTEx | STARNET | GTEx | STARNET | GTEx | STARNET | GTEx |
| **Samples** | 387 | 444 | 162 | 219 | 362 | 370 | 147 | 171 |
| **Age** | 64.3 ± 9.1 | 53.0 ± 12.8 | 69.2 ± 7.0 | 52.5 ± 12.7 | 64.4 ± 9.0 | 53.2 ± 12.8 | 69.2 ± 6.7 | 51.2 ± 12.6 |
| **BMI** | 28.7 ± 4.3 | 27.7 ± 4.1 | 29.5 ± 4.8 | 26.6 ± 4.2 | 28.7 ± 4.2 | 27.5 ± 4.0 | 29.5 ± 4.8 | 26.9 ± 4.2 |

Ideally we would probe gene-gene interactions at a genome-wide level, but Bayesian network construction is computationally intensive, and therefore, we limited this analysis to a subset of <10,000 genes that are more likely to regulate body fat distribution (Figure 1). We prioritized putative regulators of body fat distribution using three strategies: (1.) genes whose expression are co-expressed with others in adipose tissue, (2.) genes proximal to body fat distribution GWAS loci^41^, and (3.) genes that are putatively regulated by the transcription factor KLF14^23^, see Methods. For a gene to be connected to others in a co-expression network, it must be expressed in the measured dataset, must vary between samples, and must be correlated with the expression of other genes. These properties are optimal for Bayesian network construction and can indicate gene function in the tissue of interest, therefore, we constructed adipose tissue co-expression networks for all eight datasets and identified genes connected in the corresponding STARNET and GTEx networks. The union set of replicated connected genes from co-expression networks contained 7,928 genes and made up the bulk of the input to Bayesian network construction. Genes nearest to WHRadjBMI GWAS loci likely contain a mix of 30% false positives and 70% true drivers of fat distribution^71^, so we added 443 genes proximal to the 346 significant WHRadjBMI loci to the input geneset (Methods)^41,47,48^. Two previous studies identified high confidence WHRadjBMI GWAS candidate genes using colocalization methods, so we also included these 59 genes. In total, we considered this combined set of 495 genes as WHRadjBMI GWAS genes in this study. While this set does not contain all possible causal genes, it is likely enriched for them. Finally, we have previously demonstrated that *KLF14* expression regulates fat distribution in both female mice and humans^24^ and is associated with the trans-regulation of 385 genes in adipose tissue specifically^23^. We hypothesized that *KLF14*’s effect on fat distribution is mediated by some of the genes it regulates transcriptionally, and we included 385 KLF14 putative target genes in the input geneset. The union set contained 8,492 genes that were used to construct all eight Bayesian networks (Supplemental Table 1). From this diverse geneset, we aim to prioritize putative candidate GWAS genes and genes outside of GWAS loci that may play a causal role for body fat distribution.

We chose RIMBANET to construct Bayesian networks for its ability to handle large input genesets and its reproducibility between datasets^72,73^. We also added prior information about some genes to improve network performance, including genes with eQTLs in the corresponding adipose tissue depot (Supplemental Table 2). We constructed eight sex- and depot-specific networks (Methods) that contained an average of 6,250 genes connected, with an average of 6,821 directed edges between those genes (Table 2, Supplemental Table 3). Each network displayed scale-free and small-world properties consistent with known biological networks (Methods, Supplemental Table 4).

**Table 2:**
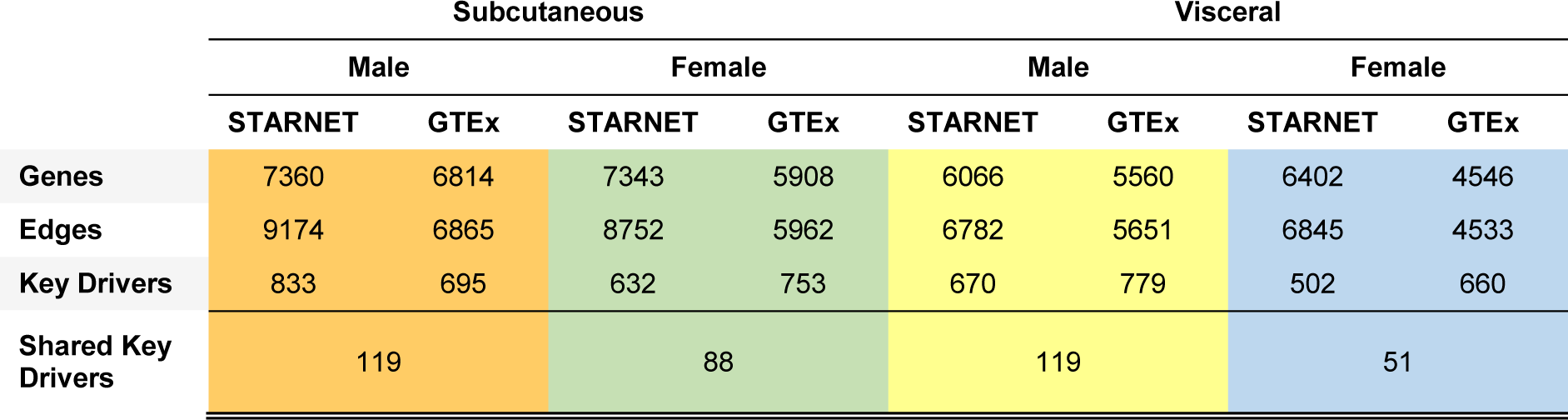
Sex- and depot-specific adipose tissue Bayesian network construction and key driver analysis results.

In general, the male networks had more connected genes and edges than the female networks, which could be due to the difference in input sample size. To test this, we randomly subsampled the male input genesets to include the same number of donors as the female networks, and constructed networks from these smaller sets. We identified an intermediate number of genes and edges for most subsampled networks (Supplemental Table 5). The sparsity of the female networks appears to be partially due to fewer female donors than male donors in the datasets.

### Bayesian Network structure identifies putative sex- and depot-specific “Key Drivers” of adipose tissue function and disease

Using the eight constructed networks, we identified putative regulators of body fat distribution, termed key driver genes (Methods, Figure 1). These are genes that regulate many genes that are part of a biological pathway or are associated with a disease or trait. Since we know that many body fat distribution genes are expressed and regulated in adipose tissue, a key driver gene that regulates many other genes in adipose tissue networks may also be a regulator of body fat distribution. Key driver genes have been biologically validated^55,58^ for their regulatory roles in Bayesian networks.

We identified an average of 691 key driver genes per network (Table 2, Supplemental Table 6). Bayesian networks, like other models, are subject to overfitting and false positive predictions, but others observe that key driver genes are more likely than other network features to be reproduced between datasets^72^ and may represent true biology. We compared the key driver predictions between our corresponding GTEx and STARNET networks and identified 334 replicated key driver genes in total (Table2, Supplemental Table 6,7). Only 38 replicated key driver genes were found in multiple sex-depot groups, and only one key driver, *ARHGEF12,* was identified in all eight networks. There were more shared key driver genes found in male networks than in female networks, and in the subsampled male networks referenced above, we identified an intermediate number of shared key driver genes, showing that the number of replicated key drivers is partially a result of the input sample size (Supplemental Table 5).

### Prioritization pipeline identifies 53 novel and putatively functional adipocyte and pre-adipocyte key driver genes

While this set of 334 replicated key driver genes likely contains many novel mechanistic drivers of fat distribution or fat storage, it also likely contains false positives and well-characterized genes. Further, we hypothesize that fat distribution is driven, in part, by adipocyte expansion, and while genes involved in other adipose tissue processes, such as tissue structure, immune function, vascularization, etc, might contribute to fat distribution or its comorbidities, these were not the focus of this study. We employed three steps to narrow this list to likely functional, testable adipocyte key driver genes (Figure 2A, Methods). First, using a curated set of six publicly available single-cell and single-nucleus RNA-sequencing studies from human and mouse adipose tissue, we identified the cell types in which each key driver gene was expressed^74–80^ and removed 110 genes that were primarily expressed in non (pre-)adipocyte cell types (Figure 2B). These genes were primarily expressed in immune cells, smooth muscle cells, and endothelial cells (Supplemental Figure 1). Second, we identified many well-studied genes in the key driver analysis, such as *FGF1, DPP4, LRP6*, and *RXRA*. While this points to the fact that our approach can identify well-known regulators of adipocyte function, we were interested in adding to the body of literature by validating genes unstudied in (pre-)adipocytes. We performed a comprehensive search of the existing literature to identify genes with known function in adipocytes and we removed 45 key driver genes involved in adipocyte processes (Figure 2C, Supplemental Bibliography part 1). Third, we used multiple lines of genetic evidence to prioritize a subset of the remaining genes. We reasoned that the set of 495 WHRadjBMI GWAS genes likely contains many genes that play a mechanistic role in fat distribution, and we prioritized 41 WHRadjBMI GWAS genes within the 179 remaining key driver genes (Figure 2D, Supplemental Table 7). Their identity as a regulator of genes within the network and their location nearby a significant GWAS locus is strong evidence that they likely have a functional role in fat distribution. Additionally, we queried the functional role these genes play when knocked out in mouse models. Although there are differences in fat storage between mice and humans, there are many conserved pathways that point to shared genetic mechanisms and similar biological outcomes^81,82^. We found that 15 genes, when knocked out in mice, affect fat storage phenotypes, and we hypothesized that they play a similar role in human fat storage (Figure 2D). In total, we prioritize 53 key driver genes with putative roles in human fat distribution via altered adipocyte fat storage, that currently have unknown function in adipose tissue (Supplemental Table 7).

**Figure 2:**
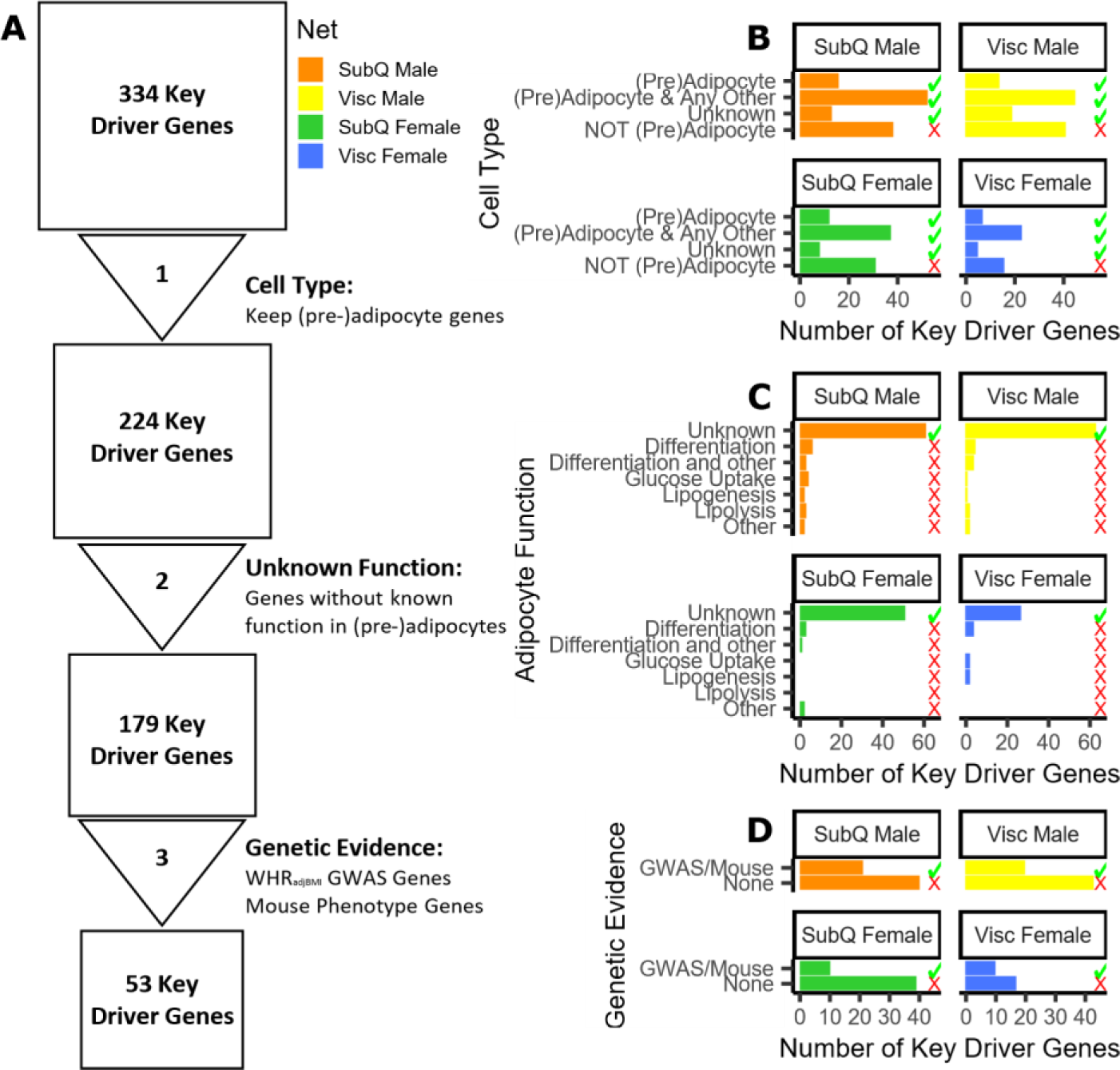
Prioritization of key driver genes for functional testing. (A) 334 key driver genes prioritized to 53 putative candidate regulators of WHR_adjBMI_ and adipocyte function using publicly available data. (B) Cell types in adipose tissue single cell- and single nucleus-RNA-seq data^74–80^ in which each key driver is expressed. (C) Key driver gene with known function in pre-adipocyte and adipocyte fat storage pathways. (D) Genetic evidence (status as WHR_adjBMI_ GWAS gene or causes mouse fat storage phenotype) for key driver genes. Green checkmark indicates genes kept in analysis pipeline, red X indicates genes removed.

We applied the same genetic evidence criteria (GWAS gene, mouse phenotype) to the 45 well-studied key driver genes that were removed in step 2 (Figure 2C). We found that almost half of these genes were identified without additional evidence from human GWAS or mouse phenotyping (Supplemental Figure 2), showing the strength of this approach as an orthogonal method with which to identify candidate functional genes.

### Four key driver genes in the Wnt signaling pathway are highly correlated with WHR adjBMI

To further characterize the 53 key driver genes, we attempted to find enrichment of specific pathway genes, but did not identify any enriched gene ontology (GO) terms or msigDB Hallmark pathways, likely due to the removal of well-characterized genes (Supplemental Figure 3). We performed a second literature search to determine the primary function of the 53 genes in other cell types and found that 13 affect the activity of the Wnt signaling pathway in other cell types (Table 3, Supplemental Bibliography part 2). These 13 genes were identified as key driver genes in all four sex-depot networks, and ten are WHRadjBMI GWAS candidate genes.

**Table 3:** Key driver genes that act in Wnt signaling in other cell types.

| Key Driver Gene | Network |  | Public Data |  |
| --- | --- | --- | --- | --- |
|  | Depot | Sex | Cell Type | Evidence |
| *BAZ1B* | Subcutaneous | Male | Multiple | GWAS |
| *HELZ* | Subcutaneous | Male | Multiple | GWAS |
| *MTMR9* | Subcutaneous | Male | Unknown | GWAS |
| *TYRO3* | Subcutaneous | Male | Unknown | GWAS |
| *ANTXR1* | Visceral | Male | Multiple | GWAS |
| *ARFGEF2* | Visceral | Male | Multiple | GWAS |
| *ARMCX3* | Visceral | Male | Multiple | Mouse |
| *ANAPC2* | Subcutaneous | Both | Adipocyte | GWAS |
| *BNIP2* | Subcutaneous | Both | Multiple | Mouse |
| *KIAA1522* | Visceral | Female | Unknown | GWAS |
| *PSME3* | Visceral | Female | Adipocyte | GWAS |
| *RSPO1* | Visceral | Female | Multiple | Mouse |
| *ZNF148* | Visceral | Female | Multiple | GWAS |

In canonical Wnt signaling, β-catenin and transcription factors TCF/LEF repress *PPARɣ* and *CEBPα* expression, which are necessary to initiate adipogenesis^30,33^ (Figure 3A). In other cell types, the 13 gene’s proteins interact with the canonical and non-canonical Wnt signaling pathway in a variety of ways (Figure 3A, Supplemental Bibliography part 2). For example, RSPO1 (r-spondin 1) is a known member of the Wnt signaling pathway; like other r-spondins, it prevents LRP degradation at the cell membrane^83,84^. TYRO3 (TYRO3 protein tyrosine kinase) activates Wnt signaling by upregulating AKT^85,86^. PSME3 (proteasome activator subunit 3, REG-ɣ, PA28-ɣ) is known to bind and target GSK3β for degradation, which releases sequestered β-catenin^87,88^. ANAPC2 (anaphase promoting complex 2), interacts with Disheveled to inhibit Wnt signaling^89^. Their role in adipose remains unknown.

**Figure 3:**
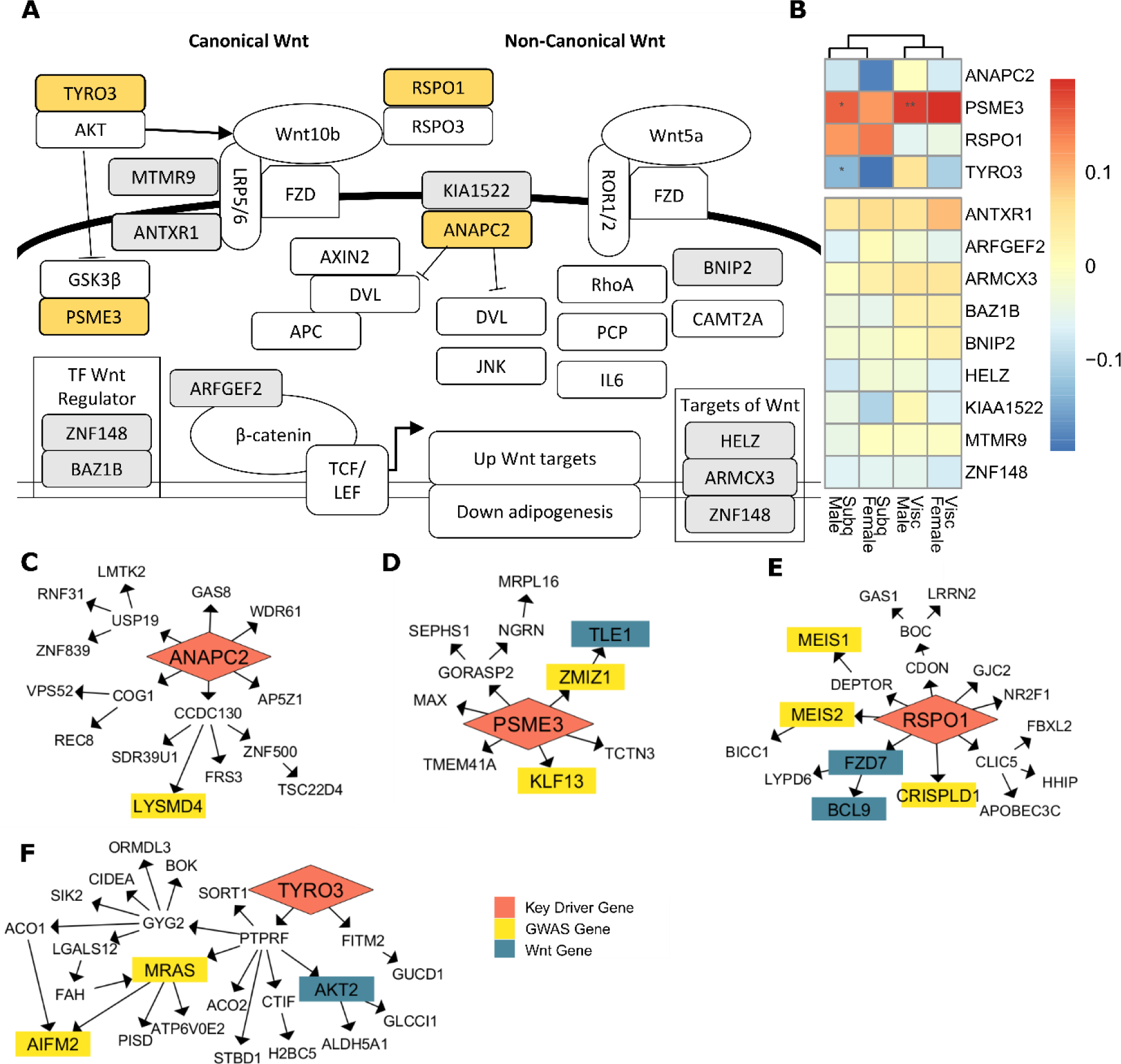
Thirteen prioritized key driver genes may affect fat storage in adipocytes through the Wnt signaling pathway. (A) The Wnt signaling pathway consists of canonical β-catenin signaling and non-canonical pathways. Key driver genes interact with Wnt pathways in other cell types (gray and yellow). (B) Key driver gene expression in adipose tissue correlations with WHR_adjBMI_ in STARNET. Pearson correlations are shown by color, p-values adjusted using FDR correction shown with * (*** = adj.P < 0.001, * = adj.P < 0.05). (C-F) Four selected key driver genes (red) regulate both WHR_adjBMI_ downstream genes^41^ (yellow) and Wnt signaling downstream genes (blue, GO term “Wnt signaling pathway”) in GTEx and STARNET. (C) ANAPC2 in the GTEX Subcutaneous Female network, (D) PSME3 in the STARNET Visceral Female network, (E) RSPO1 in the GTEx Visceral Female network, and (F) TYRO3 in the GTEx Subcutaneous Male network.

We prioritized genes related to fat distribution by calculating the correlation of each gene’s expression in each depot with overall WHRadjBMI, measured in STARNET, since GTEx did not measure WHR. We found that, while not all are significant, four genes have Pearson correlations with WHRadjBMI > 0.12 in at least one sex-depot (Figure 3B) and these correlations differ by sex or by depot. While these correlations are not numerically large because they incorporate the noise of human data and are adjusted for BMI, the > 0.12 correlations are extreme values for this data (Supplemental Figure 4). The four strongly correlated genes, *ANAPC2, PSME3, RSPO1*, and *TYRO3,* regulate a large number of downstream genes in their STARNET and GTEx networks (Figure 3 C-F). Those downstream genes contain WHRadjBMI GWAS genes (Supplemental Table 7), and often contain genes that are part of the Wnt signaling pathway. Additional evidence of involvement in fat distribution, such as expression differences between depots, WHRadjBMI GWAS signal strength, and enrichment of relevant GO terms in downstream genesets further prioritize these four genes (Supplemental Figures 5-8). *ANAPC2, PSME3,* and *TYRO3* are all found within gene dense WHRadjBMI GWAS loci, and may be the causal gene at the locus, while *RSPO1* is a putative regulator WHRadjBMI without genetic regulation.

Both subcutaneous male sub-sampled networks (Supplemental Table 5) recovered *ANAPC2* and *TYRO3* as key driver genes, showing that the subsampled networks are able to replicate the predictions of the original.

By virtue of their status as prioritized network key driver genes and their strong correlation with WHRadjBMI in humans, we hypothesize that these four genes affect fat storage in adipocytes. If the four genes also affect Wnt signaling activity in adipocytes, their effects on overall fat storage may be due to increases or decreases in Wnt signaling activity.

### ANAPC2, PSME3, and RSPO1 overexpression alter adipogenesis, not proliferation

To test these hypotheses, we first overexpressed each of the four Wnt key driver genes or a GFP control plasmid in human male pre-adipocyte cell line^90,91^ using lenti-virus (Methods, Figure 1). We were not able to perform similar experiments in female and visceral cells since no such cell lines exist at this time. We confirmed the overexpression of each gene compared to GFP controls using qPCR (Figure 4A). We also overexpressed gene *RSPO3* as a positive control, because previous literature shows that *RSPO3* has no effect on subcutaneous pre-adipocyte proliferation, but impairs adipogenesis^25^.

**Figure 4:**
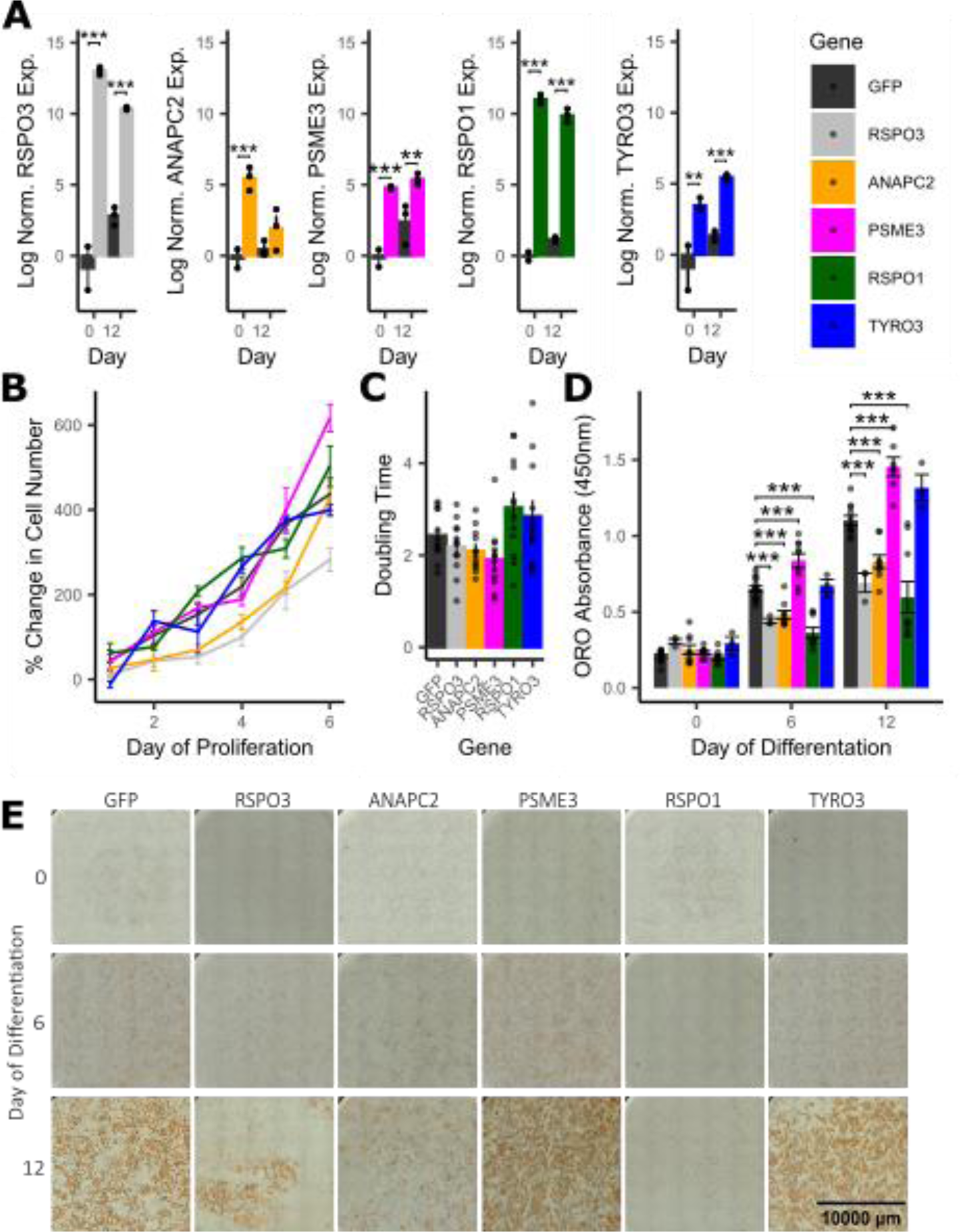
RSPO1, PSME3, and ANAPC2 affect fat storage in a pre-adipocyte cell line. (A) Expression of key driver genes compared to GFP controls at day 0 and 12 after onset of differentiation (n = 3). (B) Percent change in cell number over 6 days (n = 4). (C) Calculated doubling time in days (n = 12). (D, E) Oil Red O staining of cells was performed for each gene of interest and GFP controls at day 0, 6 and 12 after beginning differentiation, (E) representative images of one well of a 12-well plate are shown (n = 3-15). RSPO3 serves as a positive control. All plots show mean ± standard error of the mean. Differences between groups determined using 2-way ANOVA by day and gene (Gene of Interest vs GFP controls), post-hoc tests were performed using pooled t-test with Dunnett’s adjustment. Adjusted p-values shown with * (*** = adj.P < 0.001, ** = adj.P < 0.01, * = adj.P < 0.05).

We assessed the ability of these cell lines to proliferate by seeding each at the same density and counting representative wells every 24h (Methods). We found no differences in the rate of increase in cell number or the mean doubling time during exponential growth between any lines and GFP controls (Figure 4B,C).

We quantified the alterations in adipogenesis due to key driver overexpression by staining cells with the neutral lipid specific dye, Oil Red O (ORO), at 0, 6, and 12 days after the onset of differentiation. As expected, we observe an increase in lipid accumulation at day 6 and 12 compared to day 0 in all cell lines, confirming successful induction of adipogenesis, with a 4.25-fold increase in ORO absorbance in GFP controls at day 12 compared to day 0 (Figure 4D,E). Compared to GFP controls, *ANAPC2* and *RSPO1* overexpressing cells were deficient in adipogenesis, with 24.6% and 45.4% less lipid accumulation, respectively, than GFP controls at day 12. We observed that *PSME3* overexpressing cells show a 31.6% increase in lipid accumulation compared to controls at day 12. *TYRO3* overexpression caused no significant differences in lipid accumulation from controls at any time point. Positive control *RSPO3* overexpressing cells were also deficient in lipid accumulation by 37.3% compared to controls at day 12, consistent with previous studies^25^. Expression of adipocyte markers *CEBPα, PPARɣ,* and *ADIPOQ* also increased over the 12 days, and showed significant differences between *ANAPC2* and *RSPO1* overexpressing cells and controls, in agreement with adipogenesis (Supplemental Figure 9).

To fully characterize the effects of these genes, we would need to perform the same functional studies in both subcutaneous and visceral cells to determine the magnitude of adipogenic alterations in each cell type and infer effects on body fat distribution. However, pre-adipocyte cell lines from female subcutaneous tissue or any visceral tissue do not exist; therefore, in the absence of this data, we used the correlations with WHRadjBMI in STARNET (Figure 3B) to determine if our data could explain relationship between gene expression and body fat distribution. *RSPO1* findings are well aligned with human data – e.g., if RSPO1 inhibits fat storage in both subcutaneous (Figure 4D) and visceral adipocytes, then expression of *RSPO1* in visceral adipose should decrease visceral fat storage and hence decrease WHRadjBMI, and expression in subcutaneous adipose should decrease subcutaneous fat storage to increase WHRadjBMI. This is perfectly replicated in STARNET correlation data (Figure 3B) - positive correlations between subcutaneous *RSPO1* expression and WHRadjBMI, but negative correlations between visceral *RSPO1* expression and WHRadjBMI. Although *ANAPC2* and *PSME3* findings are not immediately explained by the correlation data, phenotypic effects in visceral adipocytes of different magnitude or direction could explain the observed changes.

### RSPO1 activates Wnt signaling to inhibit adipogenesis

We assessed the activity of the canonical Wnt signaling pathway using cells expressing both an overexpression plasmid and the 7TFC-luciferase construct, which measures the output of β-catenin-TCF/LEF transcription via luminescence (Methods). *PSME3* and *RSPO1* overexpressing cells were able to activate canonical Wnt signaling significantly more than GFP controls; luminescence increased by 0.74-fold and 1.34-fold in *PSME3* and *RSPO1* overexpressing cells, respectively (Figure 5A).

**Figure 5:**
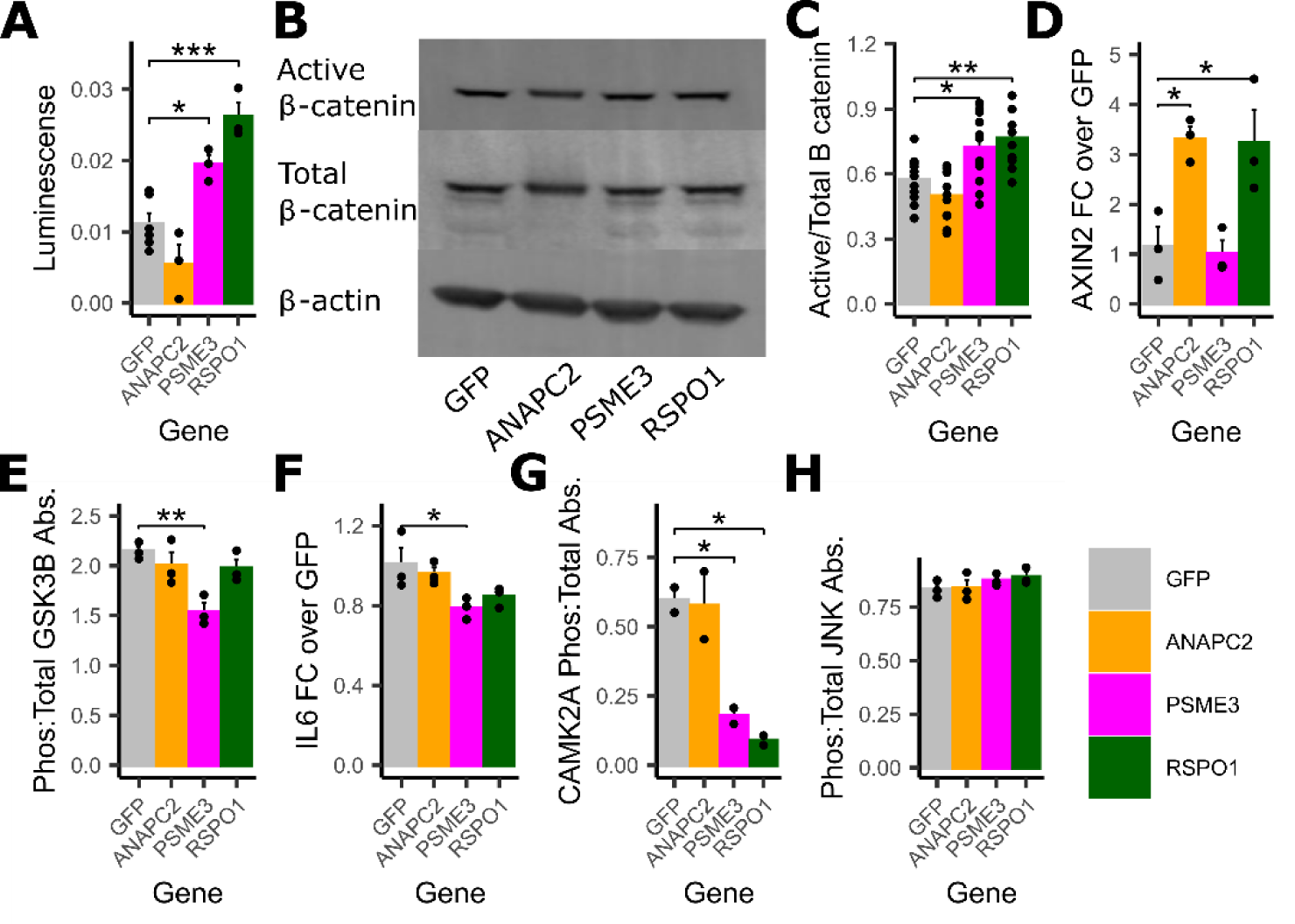
RSPO1 and PSME3 activate canonical Wnt signaling while inhibiting the Ca^2+^ non-canonical Wnt pathway. (A) Wnt transcriptional activity measured by luminescence of luciferase reporter (n = 3-6). (B) Representative images and (C) Quantification of active (non-phosphorylated) and total β-catenin by immunoblotting (n = 12). (D) Gene expression of AXIN2 measured by qPCR (n = 3). (E) Ratio of active (phosphorylated): total GSK3β measured by ELISA (n = 3). (F) Gene expression of IL6 measured by qPCR (n = 3). (G) Ratio of active (phosphorylated): total CAMK2A measured by ELISA (n = 2). (H) Ratio of active (phosphorylated): total JNK measured by ELISA (n = 3). All plots show mean ± standard error of the mean. Differences between groups determined using 1-way ANOVA by gene (Gene of Interest vs GFP controls), post-hoc tests were performed using pooled t-test with Dunnett’s adjustment. Adjusted p-values shown with * (*** = adj.P < 0.001, ** = adj.P < 0.01, * = adj.P < 0.05).

Next, we looked at individual molecules of the canonical Wnt signaling pathway^33^. We quantified the ratio of active (non-phosphorylated) β-catenin to total β-catenin using immunoblotting (Figure 5B,C), and we observed that *PSME3* and *RSPO1* overexpressing cells had a 25.4% and 33.1% increase in active/total β-catenin species compared to controls, respectively, consistent with the luciferase reporter assay. We quantified *AXIN2* mRNA expression, a target of canonical Wnt signaling, using qPCR (Figure 5D). We observed that, compared to GFP control cells, cells overexpressing *ANAPC2* and *RSPO1* increased *AXIN2* expression by 1.9-fold and 2.2-fold, respectively. Finally, using ELISAs, we measured the ratio of active to total GSK3β, an inhibitor of canonical Wnt signaling pathway (Figure 5E). Compared to GFP controls, we observed a 28.2% decrease in the ratio of active to total GSK3β in *PSME3* overexpressing cells.

We also assessed the consequences of gene overexpression on non-canonical Wnt signaling^32,92^. We observed a significant 21.7% decrease in the mRNA expression of *IL6*, a target of multiple types of non-canonical Wnt signaling, in *PSME3* overexpressing cells (Figure 5F). *RSPO1* overexpressing cells also show a non-significant 15.8 % decrease in *IL6* expression. Using ELISAs, we measured the ratio of active to total CAMK2A, a member of the Ca^2+^ non-canonical Wnt pathway, and we observed a significant 70.2% and 85.0% decrease in active to total CAMK2A ratio in *PSME3* and *RSPO1* overexpressing cells, respectively (Figure 5G). Finally, we measured the ratio of active to total JNK, a member of the planar cell polarity (PCP) non-canonical Wnt pathway and observed no differences between cell lines (Figure 5H).

### Eight key driver genes related to mitochondrial function are correlated with WHRadjBMI or *UCP1* expression

Among the 53 prioritized key driver genes, we identified a group of 13 genes that alter the function of the mitochondria in other cell types (Table 4). These genes were of particular interest, due to the variety of ways mitochondria can impact adipocyte function and lipid storage, and the 13 genes affect a variety of mitochondrial functions in other cell types (Supplemental Bibliography part 3). For example, in vascular smooth muscle cells, cardiomyocytes, and hippocampal neurons, C1QTNF3 (Complement C1q Tumor Necrosis Factor-Related Protein 3) signals through PGC-1 to increase biogenesis, oxygen consumption, and ATP synthesis^93–95^. *MIGA1’*s protein (Mitoguardin 1) is localized to the outer mitochondrial membrane (OMM) and promotes mitochondrial fusion^96,97^. *Psme3* knockout mice have larger mitochondria that are structurally dysregulated^98^ and fibroblasts overexpressing *Psme3* have elevated Bax, Cytochrome C, with concomitant anti-apoptotic effects^99^. *UBR1* (Ubiquitin Protein Ligase E3 Component N-Recognin 1) encodes an E3-ligase that degrades proteins using the N-end rule^100^. In yeast, UBR1 is localized to the OMM and targets misfolded proteins^101^ and in mouse embryonic fibroblasts, UBR1, with UBR2 and UBR4, target PINK1 in the cytosol and prevent mitophagy^102^.

**Table 4:**
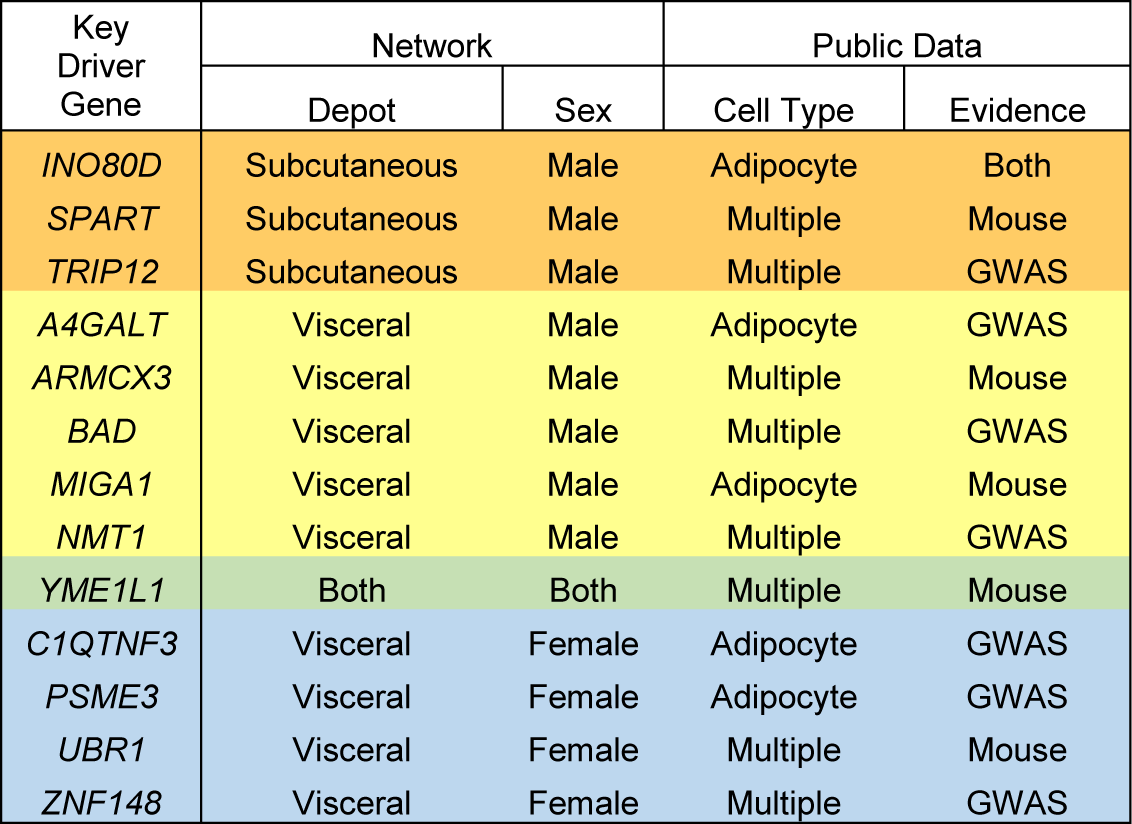
Key drivers that act in mitochondria in other cell types.

| Key Driver Gene | Network |  | Public Data |  |
| --- | --- | --- | --- | --- |
|  | Depot | Sex | Cell Type | Evidence |
| *INO80D* | Subcutaneous | Male | Adipocyte | Both |
| *SPART* | Subcutaneous | Male | Multiple | Mouse |
| *TRIP12* | Subcutaneous | Male | Multiple | GWAS |
| *A4GALT* | Visceral | Male | Adipocyte | GWAS |
| *ARMCX3* | Visceral | Male | Multiple | Mouse |
| *BAD* | Visceral | Male | Multiple | GWAS |
| *MIGA1* | Visceral | Male | Adipocyte | Mouse |
| *NMT1* | Visceral | Male | Multiple | GWAS |
| *YME1L1* | Both | Both | Multiple | Mouse |
| *C1QTNF3* | Visceral | Female | Adipocyte | GWAS |
| *PSME3* | Visceral | Female | Adipocyte | GWAS |
| *UBR1* | Visceral | Female | Multiple | Mouse |
| *ZNF148* | Visceral | Female | Multiple | GWAS |

We used depot-specific expression correlations with WHRadjBMI in STARNET to prioritize genes for further study. We identified four genes with Pearson correlation > 0.13 (Supplemental Figure 4). We also considered correlation to *UCP1* expression in STARNET, since this gene is a driver of thermogenesis in adipocytes. We found nine correlated genes, eight of which were expressed at appreciable levels (TPM ≥ 5, *INO80D* removed) in the pre-adipocyte cells used for testing (Supplemental Figure 10).

The eight mitochondrial key driver genes, including *C1QTNF3, MIGA1, UBR1,* and *PSME3,* regulate a large number of downstream genes in the corresponding STARNET and GTEx networks (Figure 6 C-F). Those downstream genes contain body fat distribution GWAS genes (Supplemental Table 7) and genes related to mitochondrial function. Additional evidence of involvement in fat distribution, such as expression differences between sexes and depots, and WHRadjBMI GWAS signal strength, further prioritized some of these genes (Supplemental Figures 6, 11-13). *C1QTNF3* and *PSME3* are found in WHRadjBMI GWAS loci along with other genes, and we hypothesize they may be the causal gene in these loci, while *MIGA1* and *UBR1* are putative mechanistic genes that are not in GWAS loci.

**Figure 6:**
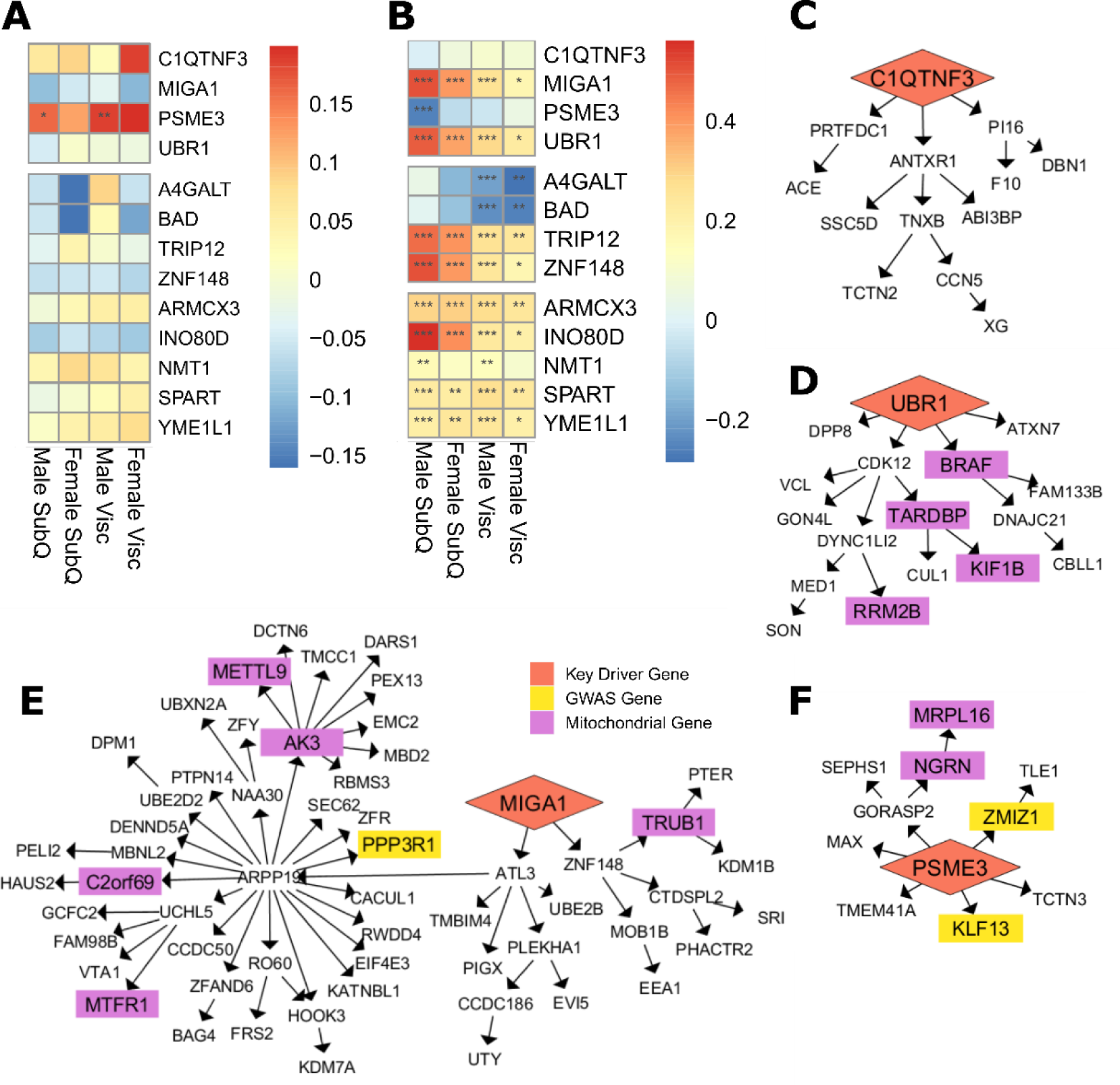
Thirteen prioritized key driver genes may affect mitochondrial function in adipocytes. (A) Key driver gene expression in adipose tissue correlations with WHR_adjBMI_ in STARNET. (B) Key driver gene expression in adipose tissue correlations with UCP1 expression in STARNET. Pearson correlations are shown by color, p-values adjusted using FDR correction shown with * (*** = adj.P < 0.001, * = adj.P < 0.05). (C-F) Four selected key driver genes are regulate both WHR_adjBMI_ downstream genes (yellow, Pulit et al) and mitochondrial downstream genes (purple, GO term “Mitochondrion”) in GTEx and STARNET. (C) C1QTNF3 in the GTEX Visceral Female network, (D) UBR1 in the GTEx Visceral Female, (E) MIGA1 in the GTEx Visceral Male network, and (F) PSME3 in the STARNET Visceral Female network.

By virtue of their status as prioritized network key driver genes and their strong correlation with WHRadjBMI in humans, we hypothesize that these eight genes affect mitochondrial function in adipocytes (Figure 1).

### Knockdown of MIGA1 and UBR1 inhibits oxygen consumption in differentiated adipocytes

To test these hypotheses, we downregulated each of the eight genes using siRNA in primary human female pre-adipocyte cells, with non-targeting siRNA as a control (Methods). We obtained mature adipocytes by differentiating these cells for 18 days, then measured the mRNA expression of each gene to confirm that the knockdown efficiency was still more than 50% compared to controls (Figure 7A). *A4GALT, ZNF148*, and *TRIP12* did not meet these criteria and *BAD* was not able to be detected, thus we removed these genes from subsequent analyses.

**Figure 7:**
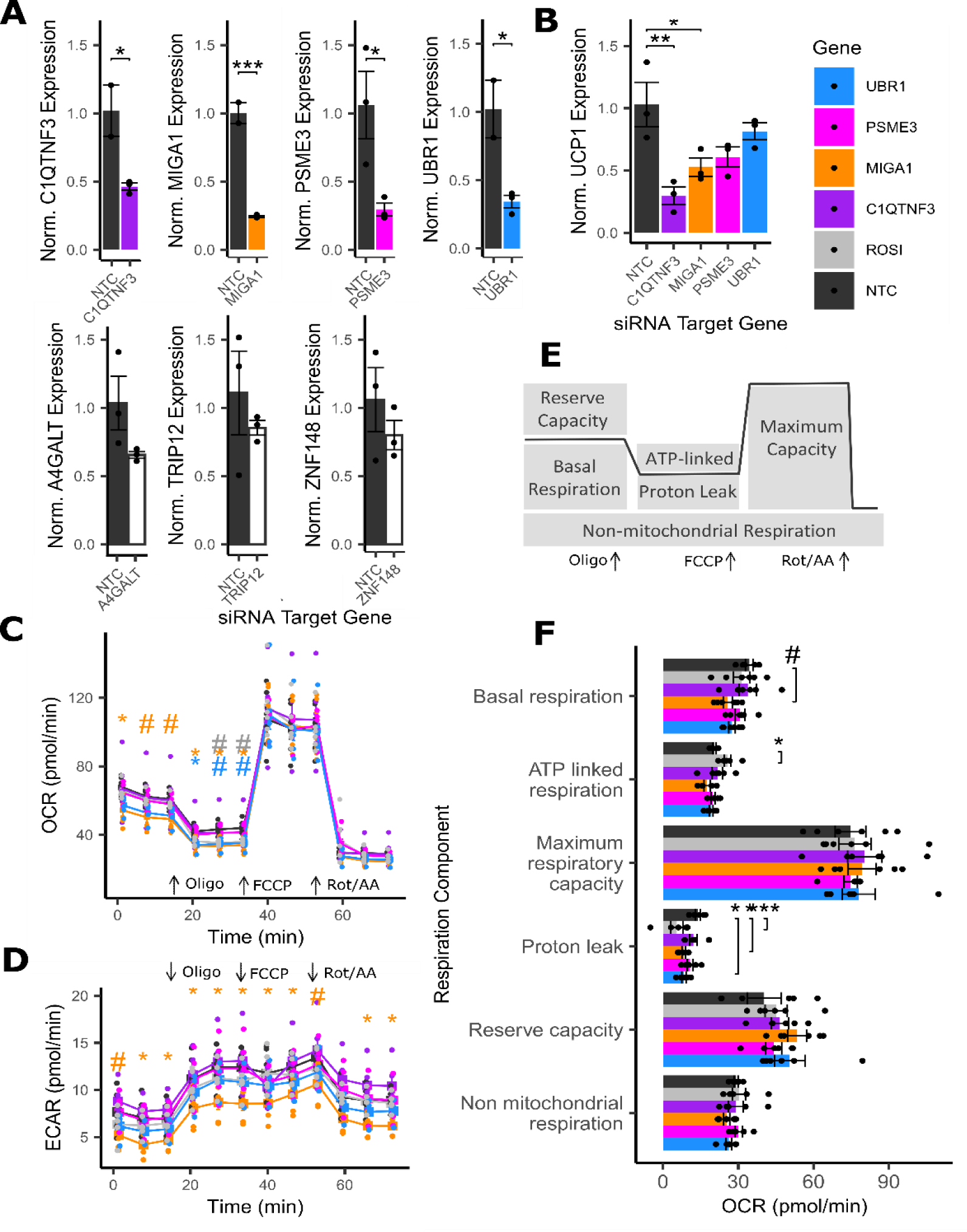
UBR1 and MIGA1 affect mitochondrial function in adipocytes. (A) Gene expression of key driver genes in non-targeting control cells and in siRNA knockdown lines. (B) (B)Expression of UCP1 in siRNA knockdown lines and controls. (C) Oxygen consumption rates genes in non-targeting control cells and in siRNA knockdown lines. (D) Extracellular acidification rates genes in non-targeting control cells and in siRNA knockdown lines. (E) Phenotypes calculated from oxygen consumption rates under various stimulation. (F) Analysis of mitochondrial phenotypes upon siRNA perturbation under stimulations. n = 6 replicates used in all assays. ROSI cells were treated with 2µM rosiglitazone for 24 h prior to assay. Differences between groups in A, B, and E were determined using 1-way ANOVA by gene (Gene of Interest vs NTC controls). Differences between groups in C and D were determined using 1-way ANOVA within each timepoint by gene (Gene of Interest vs NTCcontrols). All post-hoc tests were performed using pooled t-test with Dunnett’s adjustment. Adjusted p-values shown with * (*** = adj.P < 0.001,* = adj.P < 0.05, # = adj.P < 0.1).

We examined the effect of the knockdown of the remaining four genes on *UCP1* expression in differentiated adipocytes. We found that *C1QTNF3* and *MIGA1* knockdown significantly reduced *UCP1* expression, 71.1% and 48.7% respectively, compared to controls (Figure 7B). We also measured the expression of mature adipocyte markers. *PPARG*, *CEPBA*, and *FAPB4*. *UBR1* knockdown resulted in a significant 40.6% increase in *FABP4* expression (Supplemental Figure 14).

We then determined the effect of knockdown of each gene on cellular oxygen consumption rate (OCR) using the Seahorse assay^103^ (Methods). Importantly, we found that *MIGA1* knockdown significantly reduced the basal OCR and OCR after adding ATP-synthase inhibitor compared to controls (Figure 7C). Additionally, we found that *UBR1* knockdown significantly reduced the OCR after adding ATP synthase inhibitor compared to controls. Only *MIGA1* knockdown resulted in significantly reduced extracellular acidification rate (ECAR) in differentiated adipocytes compared to controls (Figure 7D).

Since each stimulation used in the OCR assay inhibits specific parts of the respiratory chain, we derived deeper mitochondrial phenotypes^104,105^ (Figure 7E, Methods). For example, we calculated the amount of oxygen consumed by mitochondrial proton leak, which refers to the futile H^+^ shuttling across the OMM that does not produce ATP; thermogenic uncoupling through UCP1 contributes to proton leak^106^. Both *MIGA1* and *UBR1* knockdown cells showed significantly less proton leak compared to controls; proton leak was decreased by 40.4% and 39.9% respectively (Figure 7F).

## Discussion

We present the first large-scale investigation into gene regulators of body fat distribution in adipose tissue in a sex- and depot-specific manner. We constructed large Bayesian networks using male and female adipose tissue gene expression from subcutaneous and visceral samples and identified over 300 putative regulators spanning both sexes and depots. Using additional evidence, we prioritized 53 unstudied key driver genes that may affect adipocyte function, and putatively regulate body fat distribution. Because of the unbiased nature of our initial key driver selection, we were able to prioritize putative candidate GWAS genes, as well as putative causal genes that are not in GWAS loci.

Identifying the causal GWAS gene at a given locus is a difficult task^69^, given that there can be multiple genes and signals in the locus, and the strengths of these signals can vary between populations and with biological confounders, such as sex and age. Further, while studies have found that the nearest gene to the GWAS signal is the causal gene in 70% of loci^71^, in the other 30% of loci, this assumption does not hold. For example, while *RSPO3* and *KLF14* are the only gene in the locus, the locus containing *TBX15* also contains *WARS2*, and therefore mechanistic studies had to be performed to show that *TBX15* was the causal gene^27,41^. Finally, genes not in significant GWAS loci may also contribute to the studied phenotype; in fact in the latest WHRadjBMI GWAS, SHOX2 and LRP5 are not located in significant loci, and still alter fat distribution in humans^26,29,41^.

Status as a replicated key driver provided independent evidence that allowed us to prioritize 41 of the 495 genes in WHRadjBMI GWAS loci, including *COL8A1*, *PSME3,* and *ANAPC2*. We showed here that *PSME3* and *ANAPC2* have novel functional effects on adipogenesis; *COL8A1* was also prioritized in the colocalization studies by Raulerson et al^47^ and is likely a functional regulator of fat distribution as well.

Our network analyses were able to capture known biology, as well as make novel predictions. We identified 45 key driver genes that were already well-studied in adipocytes, including 12 genes in WHRadjBMI GWAS loci and 33 putative causal genes. Most of the well-studied genes affect adipocyte fat storage by modulating adipogenesis, while a smaller number affect thermogenesis or lipolysis. *ACO1, ACAT1,* and *SLC25A1* have well-characterized effects in adipogenesis and lipogenesis (Supplemental Bibliography part 1) and are also regulated in *trans* by KLF14; these three key driver genes may mediate KLF14’s effects on female fat distribution. Our analyses also highlighted 23 novel genes in two well-established pathways, Wnt signaling and mitochondrial function, as putative drivers of adipocyte function; we demonstrated a functional role for five of the genes in these pathways. Two genes, *PSME3* and *RSPO1,* showed antagonistic effects on Ca^2+^ non-canonical and canonical Wnt signaling, consistent with literature^32^. Although the implications in adipocytes are not fully known and warrant further study, non-canonical Wnt ligands cause inflammation and vascular disease^32^ and are released from visceral fat more than subcutaneous fat^34^. While these two pathways were the focus of this paper, other pathways that contribute to fat distribution are likely represented within the 53 genes. Three validated key driver genes, *UBR1, ANAPC2*, and *PSME3,* have known roles in protein degradation; protein quality control may also be an important pathway regulating adipocyte function. Because of the bulk nature of the adipose tissue gene expression datasets, we also identified key drivers that likely act in immune cells, endothelial cells, and smooth muscle cells (Supplemental Figure 1). Although these were not the focus of our *in vitro* validation, these cell types and genes may contribute to the overall body fat distribution phenotype. Of course, key driver genes and other predictions made by networks are putative, and must be experimentally validated.

We showed that key driver gene *RSPO1* inhibited lipid accumulation by upregulating the canonical Wnt signaling pathway. This is similar to the role played by *RSPO3*, which has been shown to have different effects in visceral and subcutaneous cells that explain its effects on fat distribution^25^. In addition, *RSPO2* was recently shown to inhibit adipogenesis^84^ and *RSPO1* is a novel serum marker of obesity^107^. A recent study shows it may have a role in adipocyte beiging as well^108^. In single-cell and single-nucleus studies, *RSPO1* is expressed in pre-adipocytes and in mesothelium, the cells lining the outer wall of visceral adipose, and in GTEx, *RSPO1* expression is higher in visceral compared to subcutaneous samples (Supplemental Figure 5). RSPO1 may be released primarily from visceral mesothelial cells, upregulating Wnt to a greater degree in visceral adipose than in subcutaneous depots, which could contribute to overall differences in WHR.

We showed that key driver gene *ANAPC2* inhibited adipogenesis. Further, we saw that *ANAPC2* overexpression led to elevated *AXIN2* expression but no increases in other read-outs of Wnt signaling activity. This may explain the smaller decrease in adipogenesis compared to *RSPO1* overexpressing cells, which had a large effect on Wnt activity, although the loss of overexpression at day 12 may also account for the partial impairment of adipogenesis. *ANAPC2* encodes the cullin-like member of the anaphase promoting complex/cyclosome (APC/C) E3-ligase^109^, which degrades cell cycle machinery to promote progression to the next cell cycle phase. Interestingly, we saw no differences in proliferation due to *ANAPC2* overexpression. Studies show the necessity of exiting the cell cycle before beginning adipogenesis^110^. Others have shown that manipulating the expression of APC/C interactors *Bubr1* and *Cdc20* in cells has directionally consistent effects on adipogenesis^111,112^. *ANAPC2* may also delay and inhibit adipogenesis by continuing cell cycle mechanisms. Future experimentation is required to test this hypothesis.

We showed that key driver gene *PSME3* promoted adipogenesis, promoted Wnt signaling activity, and had no effect on mitochondrial function. Its protein product functions as a cap for the 20S proteasome^113^, and in this context REG-ɣ degrades GSK3β^88^. We confirmed that *PSME3* overexpressing cells had lower levels of active:total GSK3β than controls, although Wnt signaling is unlikely the mechanism by which *PSME3* increases adipogenesis, due to the directionally inconsistent effects. 20S proteasome member Psmb4 promotes adipogenesis in brown adipocytes by maintaining proteostasis^114^ and mutations in *PSMB8* cause lipodystrophy in humans^115^. *PSME3* has diverse roles in many cell types, with and without the 20S proteasome, including in development, fertility, cancer, and metabolism^113^; aging^116^, ribogenesis and autophagy^113^, and in lipid accumulation in hepatocytes and mouse models via sirtuin degradation^117^. Therefore, further experimentation is required to determine the mechanism by which *PSME3* increases adipogenesis.

We showed that key driver gene *MIGA1* knockdown decreased multiple components of mitochondrial function in differentiated adipocytes, suggesting that *MIGA1* contributes to normal mitochondrial function and may promote thermogenesis. Other outer membrane proteins that regulate mitochondrial fusion, MIGA2 and MFN2, also control the membrane’s interactions with lipid droplets, which affects lipid storage^118–120^. MIGA1 could affect fat storage in adipocytes through a dual role in controlling the OMM’s interactions with other mitochondria and with the lipid droplet, but further studies are required to test these hypotheses.

We showed that key driver gene *UBR1* knockdown lowered mitochondrial function in differentiated adipocytes. It had no effect on *UCP1* expression, so its role in regulating mitochondrial function is likely non-thermogenic. *UBR1* could function by maintaining protein quality and decreasing PINK1-induced mitophagy, although future studies are required to determine its precise role in mitochondria and adipocyte fat storage.

Fat distribution is a complex, full-body phenotype that is difficult to recapitulate in *in vitro* and *in vivo* models. We used subcutaneous and visceral gene expression data to make predictions, but we tested those predictions in a subcutaneous pre-adipocyte cell line that was available to us. Only *ANAPC2* was tested in the same sex-and depot-derived cell as the network in which it was a key driver. Remarkably, we observed that many predicted visceral fat key drivers had function in subcutaneous adipocyte cells, but due to these limitations, we have not fully characterized the effects of these genes on adipocyte function or fat distribution. All five genes require further studies via comparative experiments in adipocytes from subcutaneous and visceral depots. Further, the genes that were found to have no effect on adipogenesis or mitochondrial function in our studies may impact these processes in visceral adipocyte cells.

There is strong evidence that the validated key driver genes act in a sex-biased manner; therefore, follow-up studies must include cells of both sexes. Of the 53 prioritized key drivers, only five genes are replicated key drivers in both male and female networks; highlighting the unique *in silico* gene regulatory structure in each sex. Four of the validated key driver genes, *MIGA1*, *ANAPC2, PSME3,* and *RSPO1,* have a role in fertility and the development of gonads or gametes^96,121–123^. Further, RSPO1 upregulates estrogen receptor *ESR1 in vitro*^124^, and *PSME3* is located in a gene-dense female-specific WHRadjBMI GWAS locus (Supplemental Figure 4). Two of the validated genes, *RSPO1* and *PSME3*, were discovered in female visceral networks and tested in male cells, while *MIGA1* was identified in male visceral networks and tested in female cells; these may have distinct roles in the opposite sex that were not investigated in this study.

We also identified sex-bias in the network construction and key driver predictions. Both cohorts included twice as many males as females, which partially accounted for the increased complexity [genes and edges (Table 2)] and putative predictive power [number of key drivers, replicated key drivers, prioritized key drivers, and well-studied key drivers (Figure 2)] of male networks compared to female networks. However, we functionally validated three female network key driver genes of the four tested; 1/3 male genes; and 1/1 as a key driver in both sexes. Although the sample size is small, this could indicate increased predictive power of female networks. These results together certainly highlight the need to include participants of both sexes in discovery data cohorts.

We showed the validity and strength of Bayesian network modeling to predict known and novel gene regulators. We provided additional evidence of the role of Wnt signaling and mitochondria in adipocyte function, and putatively in body fat distribution. Finally, we hypothesize a broader role for five genes in regulating fat distributions in humans.

## Methods

### Data

We interrogated RNA-sequencing gene expression data from subcutaneous adipose tissue and visceral abdominal adipose tissue from the Genotype-Tissue Expression project (GTEx)^68^ and the Stockholm-Tartu Atherosclerosis Reverse Network Engineering Task (STARNET)^69^. Detailed explanations of participant inclusion, data collection, sequencing, and quantification can be found at each source. Briefly, STARNET participants are people living with coronary artery disease, from whom biopsies of abdominal subcutaneous fat and abdominal visceral fat were obtained during open thorax surgery. Samples were sequenced using the Illumina HiSeq 2000 platform. GTEx biopsies of abdominal visceral fat and leg subcutaneous fat were taken from deceased donors shortly after death. Samples were sequenced using the Illumina TruSeq platform. Both datasets were obtained in transcripts per million (TPM) format.

Expression Data Processing: We first used annotation meta-data from each source to divide the data into males and females. We used *XIST* expression to confirm these assignments. Next, we used annotations from the R package bioMart for genome build hg38 to select only the protein coding genes within each dataset. We then removed genes with less than 0.1 TPM value in greater than 80% of the samples. Finally, we log transformed the gene expression values for subsequent analysis.

### Bayesian Network Theory

Bayesian Networks are a type of Directed Acyclic Graph, where the relationships between the nodes in the graph are causal, or directed. Bayesian Networks have been used to determine directed connections between genes in an unbiased manner^54–69^. Based on Bayes’ theorem, the probability that a node in the network has a certain expression value depends on the expression of its parent nodes, or the nodes upstream of it. For a gene A, with parent genes B and C, the probability that A has a particular expression value depends on the expression state of B and C:

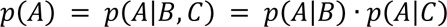

Each edge in the network represents one of these conditional relationships between genes. The total structure of the graph describes all relationships between genes as a joint probability. The joint probability of a full graph, X, is described as the geometric sum of all individual node probabilities.

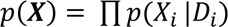

where X is the full graph, Xi is a node in the graph X, and Di is the set of parents for node Xi.

Popular methods to learn the relationships within the graph generally try to maximize a likelihood function, which increases the ability of the graph’s joint probability to describe the observed gene expression data. Since multiple graph structures can result in the same likelihood, we also add prior information to improve predictions, such as known direct connections or eQTL data.

Bayesian network construction is a time and computationally intensive process, and few popular construction tools are able to handle networks with more than 100 nodes. The Reconstructing Integrative Molecular Bayesian Network (RIMBANET)^70,125^ tool is able to handle up to 10,000 nodes by discretizing the gene expression data to reduce computational complexity. RIMBANET also takes in diverse prior information, including direct connections, eQTL data, and continuous gene expression data, to improve the predictive power of the network.

### Bayesian Network Input Genes

Co-expressed Genes: We used the python package iterativeWGCNA^126^ to obtain modules of co-expressed genes in each dataset. Weighted gene co-expression network analysis (WGCNA)^53^ employs correlations found within the data to determine which groups of genes are highly correlated and likely co-regulated. First, we computed the correlations between all genes. We raised these correlation coefficients to an empirically determined power to increase the differences observed. Next, we performed hierarchical clustering on the correlation matrix to define modules of highly correlated genes. We then assessed the success of this clustering, and iteratively reassigned genes to the modules in which they fit best. Lowly expressed or uncorrelated genes were not assigned a module. We identified which genes were assigned to modules in each of the 8 datasets. We then compared the GTEx and STARNET module assignments for each depot and sex; we found genes assigned to modules in both datasets in the 4 depot and sex groups. We then took the union set of these 4 genesets as the co-expressed geneset (Supplemental Table 1).

KLF14 *trans*-eQTL Network Genes: KLF14 predicted target genes were determined previously^23^. Single nucleotide polymorphism (SNP) rs4731702 is significantly associated with KLF14 expression in *cis* in adipose tissue of multiple cohorts^23,48^. The same variant is also associated with the expression of 385 genes across the genome in *trans.*(Supplemental Table 1).

WHRadjBMI GWAS loci-adjacent genes: The largest WHRadjBMI GWAS meta-analysis to date was performed in primarily European ancestry and discovered 346 loci associated with WHRadjBMI41. Multiple sources have determined that the functional gene is the nearest gene to the locus in ∼70% of cases^71^, so we identified genes overlapping or nearest to the lead SNP (and SNPs with LD r^2^ > 0.8) of each WHRadjBMI GWAS loci using haploReg^127^. Further, we used 2 studies that identified high quality candidate genes using colocalization methods^47,48^, where the SNPs that affect association with WHRadjBMI also affect the expression of the candidate gene, which is more likely to be functional (Supplemental Table 1).

The union set of WGCNA module genes, KLF14 targets, and putative GWAS genes made up the input to Bayesian Network construction. For each dataset, the 8,492 gene expression values were discretized into “low” “medium” and “high” bins using k-means clustering.

### Prior Information

Since multiple graph structures can result in the same likelihood score, we can use prior information to improve confidence in the network structure.

eQTLs: Following the central dogma, information flows from DNA to RNA; therefore, if SNPs influence the expression of gene A, gene A is more likely to be regulatory of other genes B and C, etc. For each dataset, we determined which genes had *cis*-eQTLs with SNPs < ± 500 kb. These eGenes were more likely to be parent nodes in the Bayesian networks. Neither STARNET nor GTEx determined *cis*-eQTLs in a sex specific manner, so these eQTL eGenes were nearly identical for male and female networks (Supplemental Table 2).

Continuous Data: Although the network is built on discretized gene expression data, RIMBANET is able to use continuous gene expression data to inform network construction. First, the continuous data is used to generate Pearson correlations between all genes. Correlations with significance p < 0.01 are used as prior information to determine possible parents and prioritize which edges to add or remove.

### Bayesian Network Construction

Bayesian Networks for each dataset were constructed using RIMBANET using the discretized gene expression data, a list of eQTL eGenes, and the continuous gene expression data. The RIMBANET shell script was adapted for implementation on the University of Virginia’s high-performance computational cluster (Rivanna). RIMBANET was run with these tags: –C TRUE to specify continuous data, -w to add the continuous dataset, -d to add the discretized dataset, -e to add eQTL eGenes. RIMBANET creates 1,000 versions of each network with random initial seeds. RIMBANET employs a Markov blanket to identify potential parents for each gene, then iteratively adds edges with the highest prior information to create the best network structure to represent the data, as measured by the Bayesian Information Criterion (BIC).

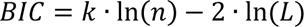

where n is the number of nodes, k is the number of parameters estimated (edges), and L is the likelihood function. Maximizing this function results in the best fit graph structure that is not overly complex. Then, RIMBANET merges the 1,000 network versions by retaining edges present in 30% of the iterations. Finally, RIMBANET produces a directed acyclic graph by removing complete cycles.

### Properties of Biological Networks

Biological networks share some common features, including reproducibility, scale-free-ness, and small world-ness^128^. Scale-free networks have degree distributions that follow the power law; most nodes have a small number of downstream genes, while a few nodes have a large number of downstream genes^129^. The distribution of the number of downstream genes per parent can be fitted as a power law P(k) ∼ k^-a^, where k represents the number of downstream edges and P(k) represents the fraction of nodes that have k edges. The degree exponent, a, is determined by fitting the line log(P(k) ∼ -a log(k). In biological networks, the degree exponent is commonly 2-3. We used the igraph() package in R to calculate the degree exponent for each network. Small world networks are highly clustered, yet have a short average distance between nodes^130^. We used the qgraph() package in R to calculate the clustering coefficient and average path length between nodes for each network. Since these properties scale with the number of nodes in the network, we compared these metrics to a random graph of the same size.

### Key Driver Gene Analysis

Key driver genes of each network were identified with two methods: 1. by the number of downstream genes regulated by each potential key driver gene and 2. by the enrichment of disease genes in the set of downstream genes regulated by each potential key driver gene.

To identify type 1 key driver gene testing, first, every gene in the network was profiled to determine its number of downstream genes at distances 1-10 edges away using the shortest path. Then, the mean and standard deviation in the number of downstream genes at each edge distance was calculated for the network.

For each potential key driver gene, we calculated ten score functions:

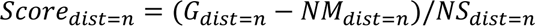

where n is the distance in edges, G is the number of downstream genes the potential key driver gene has at distance n, NM is the network mean number of downstream genes at distance n, NS is the network standard deviation in the number of downstream genes at distance n. This is a metric of the extremeness of G, effectively a z-score.

Finally, we calculated a total score for each potential key driver gene by summing the ‘z-scores’, weighting smaller edge distances away from the potential key driver higher than large edge distances:

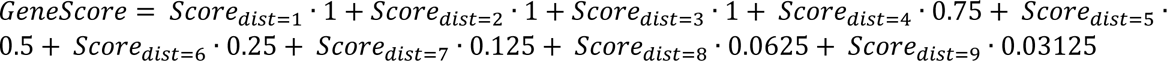

The top 10% highest scoring genes in the network were declared type 1 key driver genes, similar to previous studies^55^.

To identify type 2 key driver genes, a ‘neighborhood’ of downstream genes was declared for every gene in the network as those genes within 4 edges away. Next, WHRadjBMI GWAS genes^41^, defined above, were identified within the downstream neighborhood. Finally, using Fisher’s exact test, we determined whether the number of GWAS genes in the downstream neighborhood was significantly more than expected by chance, compared to the whole network. Genes with significant downstream enrichments for WHRadjBMI GWAS genes were declared type 2 key driver genes.

Key driver genes for each network were the union set of type 1 and type 2 key driver genes. Shared key driver genes were genes identified as either type 1 or type 2 key driver genes in both STARNET and GTEx networks of the same type.

### Testable Key Driver Gene Selection

#### Identification of cell type

We used 7 publicly available single cell- or single nucleus-adipose tissue or adipose tissue derived-stromal vascular fraction RNA sequencing datasets from both human and mouse^74–80^. Since there was some disagreement between studies, all cell types in which the gene was expressed in any study are reported in Supplemental Table 7. We removed genes that were only expressed in non-adipocyte cell types.

#### Identification of Function in Adipocytes

We identified well-studied key driver genes using a comprehensive literature search (Supplemental Bibliography part 1). For each gene, we used GeneCards to identify alternate names for each gene or corresponding protein. We then searched PubMed and Google Scholar for functional studies in cells demonstrating a role for that gene in pre-adipocytes or adipocytes. Terms searched include “adipocyte”, “adipogenesis”, “differentiation”, “lipogenesis”, “lipolysis”, “glucose uptake”, “browning”, and “thermogenesis”.

#### Additional Genetic Evidence

We prioritized key driver genes based on two types of genetic evidence. First, we identified WHRadjBMI GWAS genes^41^ within the set of key driver genes. Second, we prioritized genes involved in fat storage and distribution in mouse models. We queried the Mouse Genome Informatics database^131^ and the International Mouse Phenotyping Consortium^132^ to determine if the gene knockout in mice results in significant differences in fat pad size, total body fat mass, lean mass, or related phenotypes. We did not consider overall body size differences, as these may be indicative of BMI related phenotypes.

### Lentivirus Construction

Overexpression plasmids were constructed by VectorBuilder (VectorBuilder Inc, Chicago, Illinois, USA) using the mammalian gene expression lenti-viral vector backbone with 1 open reading frame. This backbone contains 3^rd^ generation lenti-viral integration sites and ampicillin resistance. Using GTEx, we identified the most abundant isoform in adipose tissue for each gene. This isoform was added to the plasmid, followed by a P2A linker, then the GFP reporter sequence. This construct was under the CMV promoter. Control plasmids contain the GFP reporter gene under the CMV promoter with no P2A linker.

7TFC was a gift from Roel Nusse, purchased from Addgene (Addgene, Watertown, Massachusetts, USA; Addgene plasmid # 24307). The 7TFC plasmid contains 3^rd^ generation lenti-virus integration sites, the Firefly Luciferase gene under 7 repeats of the TCF promoter and an mCherry marker under the SV40 promoter and ampicillin bacterial resistance gene.

We obtained the plasmids in *E. coli* swabs in agar. We cultured the *E. coli* on Luria-Bertani broth agar plates containing 100 µg/mL ampicillin, and sub-cultured single colony forming units in 50 mL Luria-Bertani broth containing 100 µg/mL ampicillin. Plasmids were isolated from *E. coli* using the Nucleobond Xtra Midi prep kit (Takara Bio, San Jose, California, USA) following the manufacturer’s protocol (Cat# 740422.50). Plasmids were packaged into 3^rd^ generation replication deficient lenti-virus in HEK-293T cells using the Lenti-Pac HIV Expression Packaging Kit (Genecopoeia, Rockville, Maryland, USA) following manufacturer’s protocol (Cat# LT001).

### Transduction and Sorting of Human Pre-adipocyte Overexpression Cell Lines

We obtained human male pre-adipocyte Simpson-Golabi-Behmel syndrome (SGBS) cells from Dr. Martin Wabitsch^90^ at passage number 35-40. All pre-adipocyte cells were grown and maintained as described previously^91^ in DMEM:F12 media (ThermoFisher Scientific, Waltham, Massachusetts, USA) containing 10% Fetal Bovine Serum, 1% Penicillin/Streptomycin, 8.1 ng/mL biotin and 3.5 ng/mL pantothenate. During transduction with lenti-virus, the fetal bovine serum was first heat-inactivated at 65 C for 30 minutes, and 8 µg/mL polybrene was added to improve transduction efficiency. We plated cells in 6 well plates and grew them to 70% confluence before transducing the cells with lenti-viral-particles containing each of the plasmids listed above. Cells containing high levels of GFP or mCherry were sorted using the FACS Aria Fusion Cell Sorter (BD Biosciences, Franklin Lakes, New Jersey, USA). We used cells with passage number less than 46 for subsequent assays.

### siRNA of Target Genes

**Table 5:**
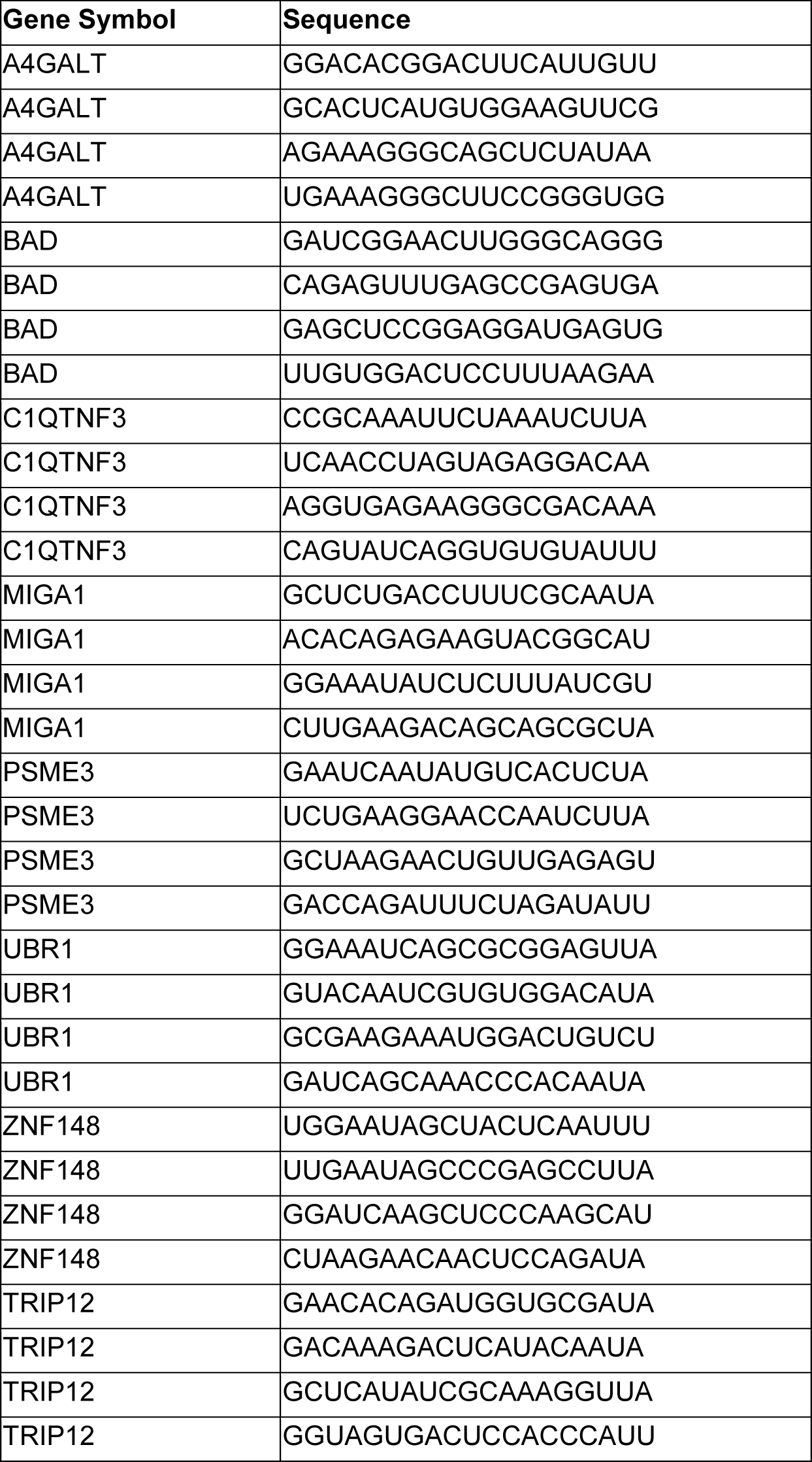
siRNA sequences.

### Transfection and Differentiation of Human Primary Pre-adipocytes

Human female subcutaneous primary pre-adipocytes were purchased from Zenbio (Zenbio, Cat# SP-F-SL; Lot# SL0061) and were differentiated according to the standard Zenbio white adipocyte differentiation protocol. Briefly, human primary subcutaneous pre-adipocytes were seeded on collagen-coated 96-well plate (20,000cell/well, Corning, 354650) with 200µl of PM-1 medium (Zenbio, #PM-1) and the cells were established overnight, then transfected with siRNAs of target genes or scramble controls for 3 days using Lipofectamine RNAiMAX (Invitrogen, cat# 13778-150). The culture medium was replaced with 150µl of differentiation medium DM-2 (Zenbio, #DM-2), and cells were cultured for 7 days. Media was replaced with maintenance medium AM-1 and cells were cultured for additional 7 days for the following qPCR and Seahorse experiments.

### Quantification of Gene Expression in lenti-viral treated Human Pre-Adipocytes and Differentiating Cells

We grew cells in 12 well plates. Once they reached confluency, they were washed with Phosphate Buffered-Saline (PBS) and incubated in 400 µL Trizol, then scraped and harvested. We extracted RNA using the RNeasy Micro Kit (Qiagen, Velno, Netherlands), following manufacturer’s protocol (Cat# 74004). We digested the DNA species on the Qiagen spin column using the RNAse-free DNAse kit (Qiagen, Velno, Netherlands) following manufacturer’s protocol (Cat# 79254). We quantified the isolated RNA using the Qubit with RNA Broad Range assay kit (ThermoFisher Scientific, Waltham, Massachusetts, USA), following manufacturer’s protocol (Cat# Q10210). We reverse transcribed cDNA from the RNA templates using SuperScript IV Reverse Transcriptase Kit (ThermoFisher Scientific, Waltham, Massachusetts, USA) with Oligo(dT)20 primers, following manufacturer’s protocol (Cat# 18090010). We quantified cDNA abundance using quantitative-polymerase chain reaction (qPCR). Samples and standard curves were prepared using GoTaq qPCR Master Mix (Promega, Madison, Wisconsin, USA) and gene specific primers (Integrated DNA Technologies, Coralville, Iowa, USA) (Table 4). Samples were measured using the QuantStudio 5 Real Time PCR system (ThermoFisher Scientific, Waltham, Massachusetts, USA), and were analyzed using Thermo Fisher Connect qPCR Standard Curve analysis software.

### Quantification of Gene Expression in siRNA treated Human Differentiated Adipocytes

The total RNA samples were isolated from human differentiated adipocytes using KingFisher™ Flex Magnetic Particle Processor according to the MagMAX™ mirVana™ Total RNA isolation protocol. We obtained the cDNA samples from the RNA templates using SuperScript IV Reverse Transcriptase Kit (ThermoFisher Scientific, Waltham, Massachusetts, USA) with Oligo(dT)20 primers, following manufacturer’s protocol (Cat# 18090010). We quantified cDNA abundance using quantitative-polymerase chain reaction (qPCR) using the QuantStudio 5 Real-Time PCR system (ThermoFisher Scientific, Waltham, Massachusetts, USA). The conditions were: 42 °C for 5 min, a 10s denaturation step at 95 °C, followed by 40 cycles of 95 °C for 5s and 58 °C for 40s.

**Table 6:**
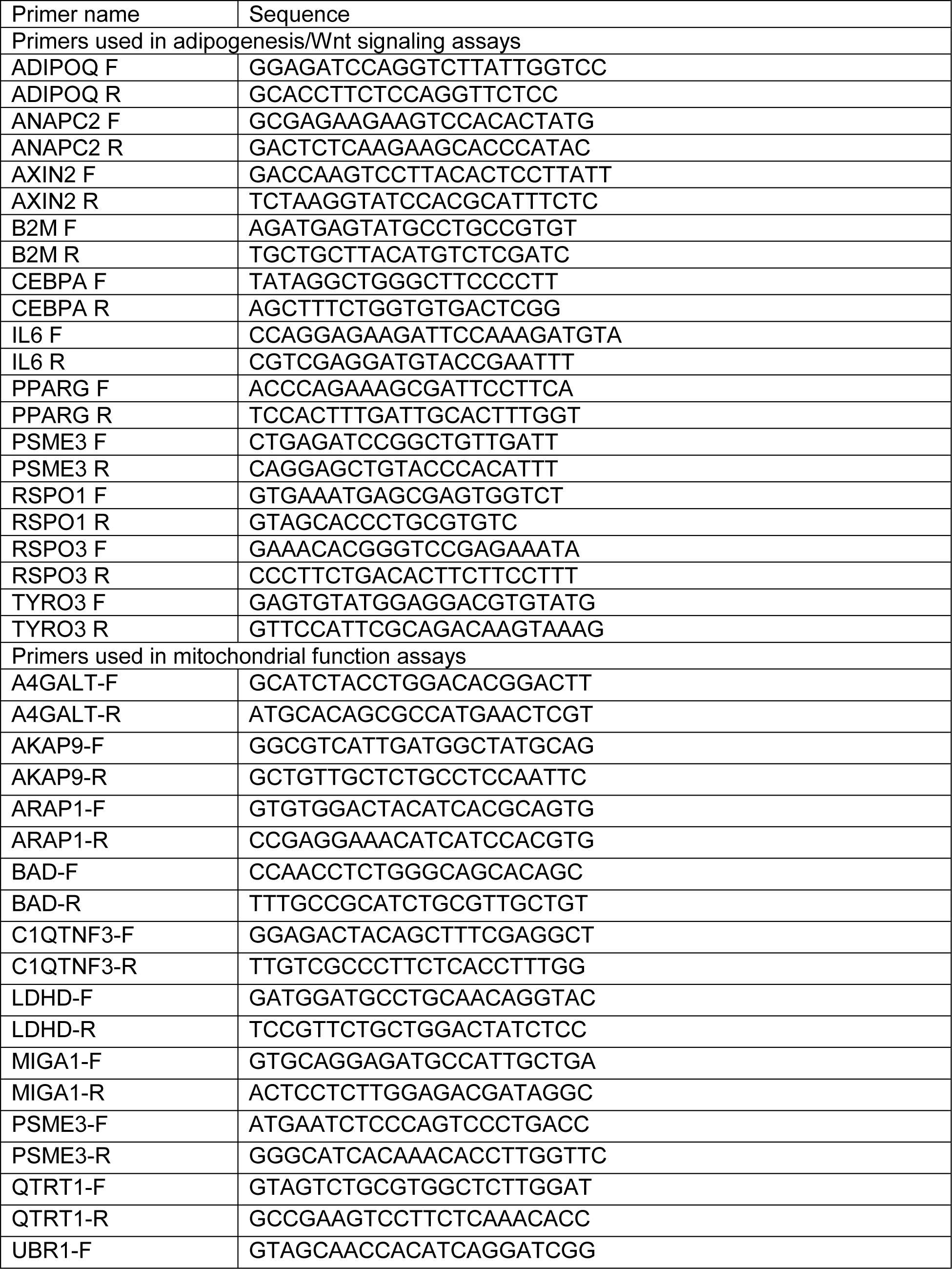
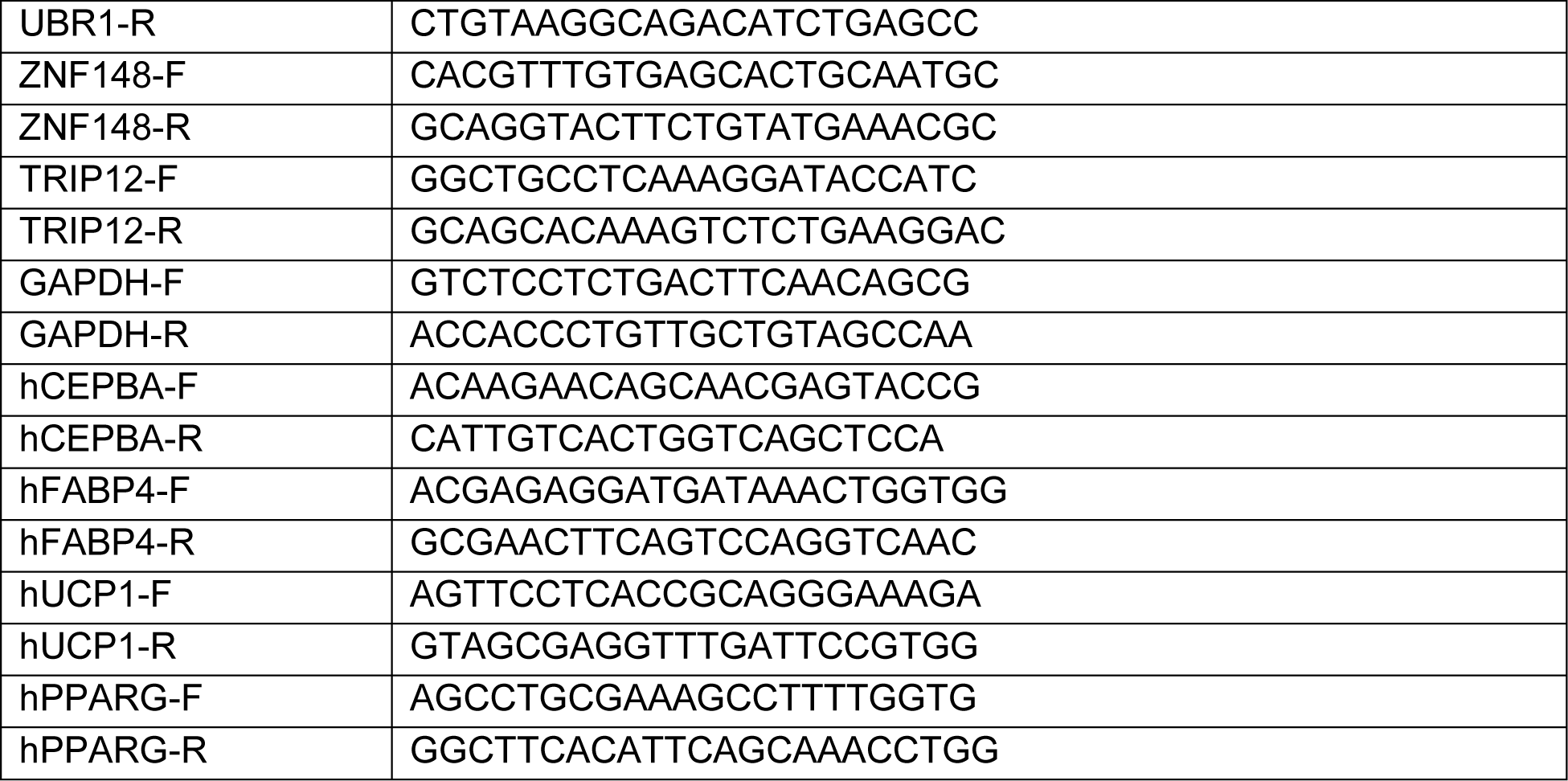
qPCR Primer Sequences.

### Cellular Phenotyping

*Proliferation Assay:* We plated 20,000 cells/well in 24 well plates. After 24 hours, 4 wells were trypsinized and cells were counted using a hemocytometer. Wells were washed with PBS and growth media was replaced. Every 24 hours, 4 more wells were counted, for a total of 6 days. Growth media was replaced every 2 days. Most cells reached exponential growth by day 4 (Figure 4B). We then calculated the doubling time of each cell line using the formula:

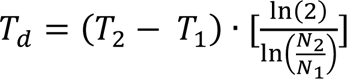

where T_d_ is doubling time, T_1_ and T_2_ are initial and final time measurements, and N_1_ and N_2_ are the initial and final quantity of cells.

*Adipogenesis Assay:* We differentiated cells into lipid-containing adipocytes as detailed previously^91^. Briefly, we plated 40,000 cells/well in 12 well plates. Cells were incubated for 2-5 days until they reached 100% confluency, then incubated for 48 hours post-confluency. Adipogenic media (DMEM:F12, 1% Penicillin/Streptomycin, 8.1 ng/mL biotin, 3.5 ng/mL pantothenate, 0.01 mg/ml transferrin, 20 nM insulin, 100 nM cortisol, 0.2 nM triiodothyronine, 25 nM dexamethasone, 250 µM 3-isobutyl-1-methylxanthine, and 2 µM rosiglitazone) was added to each well to initiate differentiation. After 4 days, we changed the media to DMEM:F12, 1% Penicillin/Streptomycin, 8.1 ng/mL biotin, 3.5 ng/mL pantothenate, 0.01 mg/ml transferrin, 20 nM insulin, 100 nM cortisol, and 0.2 nM triiodothyronine. Every 4 days, this media was replaced.

*Quantification of Adipogenesis:* We quantified the amount of lipid stored in cells using Oil Red O (ORO) dye. 0.25 grams of dye was suspended in 48 mL of 98% isopropanol and 32 mL of DI water. Unsuspended dye was removed from the ORO solution using 0.045 µM vacuum filtration. We repeated filtration (∼3X) every 24 hours until no precipitate was observed. Cells were washed with PBS, then fixed in 300 µL 4% paraformaldehyde for 15 minutes. Cells were washed with 60% isopropanol, then dried completely. 250 µL of ORO solution was added to the cells for 5 minutes. Cells were washed with DI water twice, then dried completely. We imaged the full wells using the EVOS microscope (below). Oil Red O dye was then eluted from cells in 200 µl of 100% isopropanol for 2 minutes. Eluted ORO was quantified by measuring absorbance at 450 nm.

*Imaging:* We took images using the EVOS M7000 imaging system (ThermoFisher Scientific, Waltham, Massachusetts, USA) at 10x magnification, using phase contrast and color. We constructed full well composite images by taking 30 adjacent images in a 5x6 grid that covers most of the well. Composite images were stitched together using imageJ:Fiji plugin Grid/Collection Stitching^133^.

### Quantification of Wnt Signaling

*Quantification of Wnt Signaling Transcriptional Activation:* We performed luciferase assays using the SGBS:7TFC reporter line. SGBS:7TFC cells were transduced with lenti-virus containing the gene of interest or GFP control plasmids. Images were taken to ensure a high percentage of dual mCherry and GFP expressing cells. 10,000 cells/well were plated in clear bottom, white-walled 96 well plates, with 6 replicates of each gene or control. After 24 hours of incubation, luciferase activity was measured using the Luciferase Assay System (Promega, Madison, Wisconsin, USA) following manufacturers protocol (Cat# E1500). Briefly, the cells were lysed in 20 µl lysis buffer, 100 µl of luciferin-containing reagent was added, then emitted light was measured for 10 seconds using a luminescence plate reader. Luminescence readouts in each well were normalized to mCherry fluorescence to account for total luciferase insertions by the 7TFC cassette.

*Quantification of Protein Activation:* Active (Ser33/Ser37/Thr41 non-phosphorylated) β-catenin and total β-catenin species were measured using western blotting. Total proteins were isolated in RIPA buffer containing 1% protease and 1% phosphatase inhibitors (ThermoFisher Scientific, Waltham, Massachusetts, USA, Cat# 89901, Cat# 78429, Cat# 78426). We quantified total protein species using the bicinchoninic acid (BCA) assay (ThermoFisher Scientific, Waltham, Massachusetts, USA) following the manufacturer’s protocol (Cat# 23225). We denatured samples at 70°C for ten minutes, then ran 20 µg total protein on a NuPAGE 4-10% BisTris Gel at 240V for 40 minutes (ThermoFisher Scientific, Waltham, Massachusetts, USA, Cat# NP0336BOX). We transferred the protein to an Immobilon-FL PVDF membrane at 80V for 60 minutes (MilliporeSigma, Burlington, Massachusetts, USA, Cat# IPFL00010). We labeled active and total β-catenin and β-actin control bands using primary antibodies (Cell Signaling Technologies, Danvers, Massachusetts, USA; Non-phospho (Active) β-Catenin (Ser33/37/Thr41) (D13A1) Rabbit mAb Cat#8814, dilution 1:500; β-Catenin (15B8) Mouse mAb Cat#37447, dilution 1:1000). Bands were labelled with fluorescently conjugated secondary antibodies (ThermoFisher Scientific, Waltham, Massachusetts, USA; Goat anti-Mouse IgG (H+L) Cross-Adsorbed Secondary Antibody, Cyanine3, Cat# A10521, dilution 1:20,000; Goat anti-Rabbit IgG (H+L) Cross-Adsorbed Secondary Antibody, Cyanine3, Cat# A10520, dilution 1:20,000). We imaged the labelled protein on Amersham Imager 600 (Global Life Sciences Solutions, Marlborough, Massachussets, USA) using RGB fluorescence settings. Densitometry calculations were performed using imageJ.

Amount of active and total GSK3β, JNK, and CAMK2A were quantified using Enzyme-linked immunoassays (ELISA). Cells were harvested and lysed according to each manufacturer’s protocol. Active GSK3β (Ser9 phosphorylated) and total GSK3β were measured using an ELISA kit (RayBiotech, Peachtree Corners, Georgia, USA) using manufacturer’s protocols (Cat# PEL-GSK3b-S9-T). Active JNK (Thr183/Tyr185 phosphorylated) and total JNK were measured using an ELISA kit (RayBiotech, Peachtree Corners, Georgia, USA) using manufacturer’s protocols (Cat# PEL-JNK-T183-T-1). Active CAMK2A (Thr286 phosphorylated) and total CAMK2A were measured using an ELISA kit (Assay BioTechnology, Fremont California, USA) using manufacturer’s protocols (Cat# FLUO-CBP1509 and CB5092).

### Quantification of oxygen-consumption rate (OCR) and extracellular acidification rate (ECAR)

Oxygen consumption rate (OCR) and extracellular acidification rate (ECAR) was determined using a Seahorse XF96 analyzer in combination with the Seahorse mitochondrial stress test kit according to a standard protocol^103^. In brief, human primary pre-adipocytes were plated and differentiated as described above. Differentiated cells were washed with DPBS twice and incubated with Seahorse XF assay medium supplemented with 2 mM glutamax, 10 mM glucose, 1 mM sodium pyruvate (PH 7.4) for 45 min at 37 °C in a non-CO2 environment. Both OCR and ECAR were subsequently measured in real time using XF96 extracellular flux analyzer (Seahorse Bioscience). The optimized concentration of compounds for mito-stress assay were 1.5 μM of oligomycin, 1.5 μM of carbonyl cyanide-p-trifluoromethoxyphenylhydrazone (FCCP), and 0.5 μM of Rotenone&antimycin A. Following the extracellular flux analysis, the OCR and ECAR were normalized by cell number, quantified using Hoechst staining.

*Quantification of deep OCR phenotypes:* We calculated basal mitochondrial respiration, ATP-linked respiration, proton leak, maximal respiratory capacity, reserve capacity, and non-mitochondrial respiration from the OCR assay as described previously^104,105^. We defined condition ‘A’ as timepoints 1, 2, and 3 under basal stimulation; condition ‘B’ as timepoints 4,5,6 under oligomycin stimulation; condition ‘C’ as timepoints 7,8,9 under FCCP stimulation; and condition ‘D’ as timepoints 10,11 and 12 under Rot/AA stimulation. We considered each of the three timepoints within each condition as technical replicates, and the six samples as biological replicates. We averaged the three timepoints per condition into one value per biological replicate. We then defined non-mitochondrial respiration as D; basal mitochondrial respiration as A - D; ATP-linked respiration as A - B, proton leak as B - D; maximal respiratory capacity as C - D; and reserve capacity as C-A.

## Statistical Methods

Differences in proliferation and differentiation assays using the GFP-expressing control cells and cells expressing the gene of interest were assessed using 2-way ANOVA by gene and time (day). Post-hoc tests were performed between GFP controls and genes of interest within each timepoint using pooled t-tests with p-value adjustment using Dunnett’s adjustment. Differences in Wnt signaling using the GFP-expressing control cells and cells expressing the gene of interest and mitochondrial assays using the non-targeting control-expressing cells and cells with siRNA for the gene of interest were assessed using 1-way ANOVA by gene. Post-hoc tests were performed between controls and genes of interest using pooled t-tests with p-value adjustment using Dunnett’s adjustment. Analyses were performed using base R’s anova() function and the emmeans package’s emmeans() and contrasts(). We reported only significant p-values, with the exception of Figure 7 where indicated. All bar plots display the mean, with error bars displaying the standard error of the mean (S.E.M).

## Supporting information

Supplemental Figures 1-14, Bibliography

## Acknowledgements

This work was supported by NIH/NIDDK grant R01 DK118287 and NIH training grant T32 HL007284.

## Data and code availability

Gene expression and eQTL data from GTEx can be found at dbGaP Accession phs000424.v8.p2on 10/01/2020. Gene expression and eQTL data STARNET can be found at https://www.ncbi.nlm.nih.gov/projects/gap/cgi-bin/study.cgi?study_id=phs001203.v3.p1, data available on request. Code used in these analyses can be found at https://github.com/jnr3hh/Reed_Civelek_2023_manuscript.

