## Supplemental Figures 1-14, Bibliography for "Systems genetics analysis of human body fat distribution genes identifies Wnt signaling and mitochondrial activity in adipocytes"

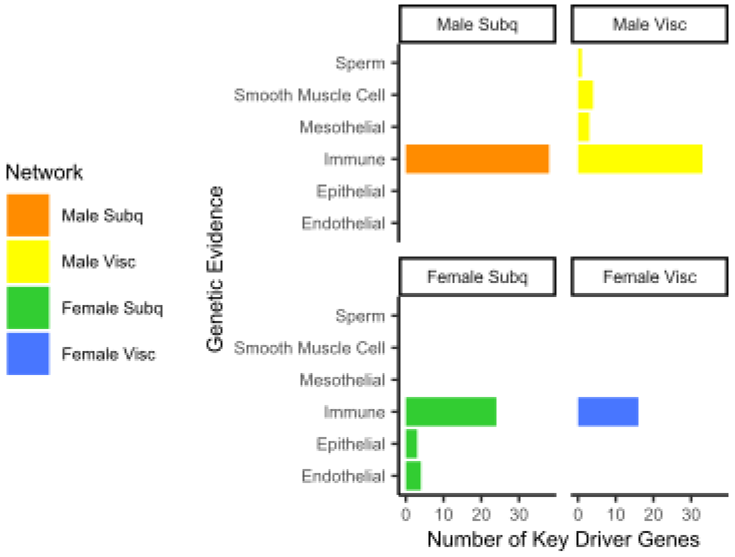


*Supplemental Figure 1: The 110 key driver genes removed from analyses due to primary expression in other cell types, in Figure 2, step 1.*

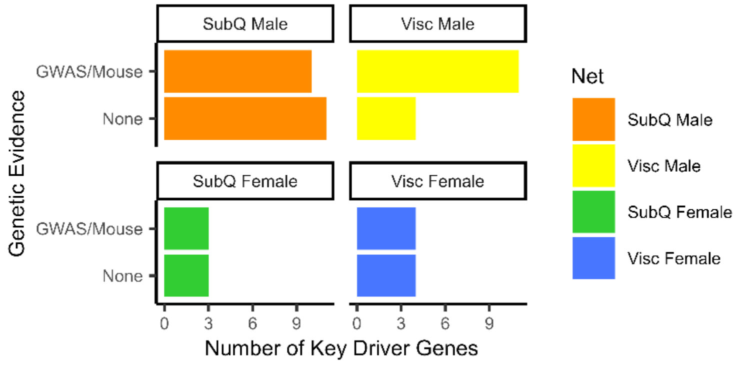


*Supplemental Figure 2: The 45 key driver genes with known function in adipose tissue that were removed in Figure 2, step 2 were subjected to the Genetic Evidence criteria used in Figure 2, step 3.*

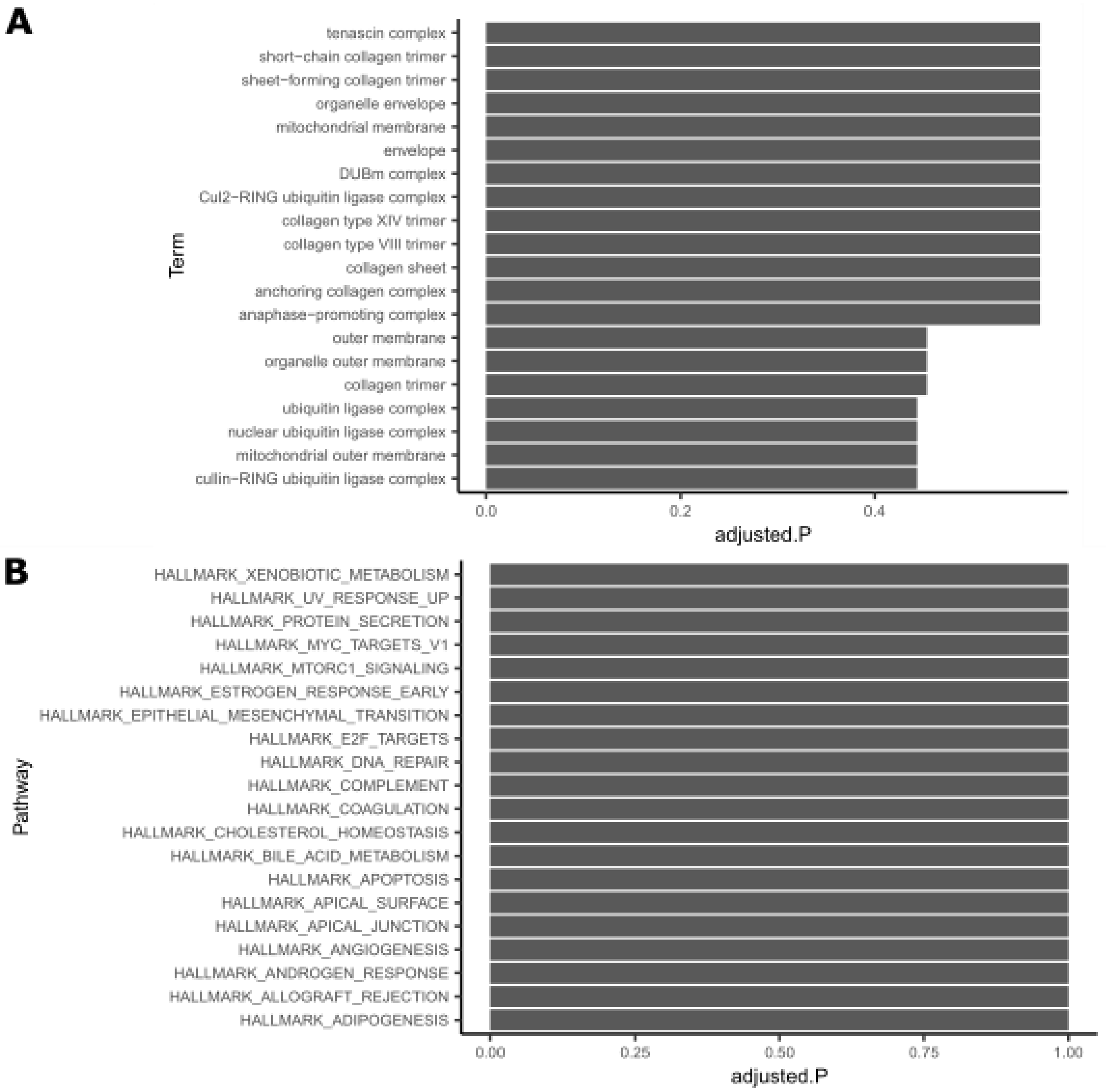


*Supplemental Figure 3: The 53 key driver genes prioritized for further study were not enriched for any specific pathways. (A) The top 20 Gene Ontology (GO) biological processes pathways, tanked by adjusted p-value. (B) The top 20 msigDB Hallmark pathways, ranked by p-value, adjusted p-value shown. Fisher's exact test used to test enrichments, FDR correction used to adjust p-values.*

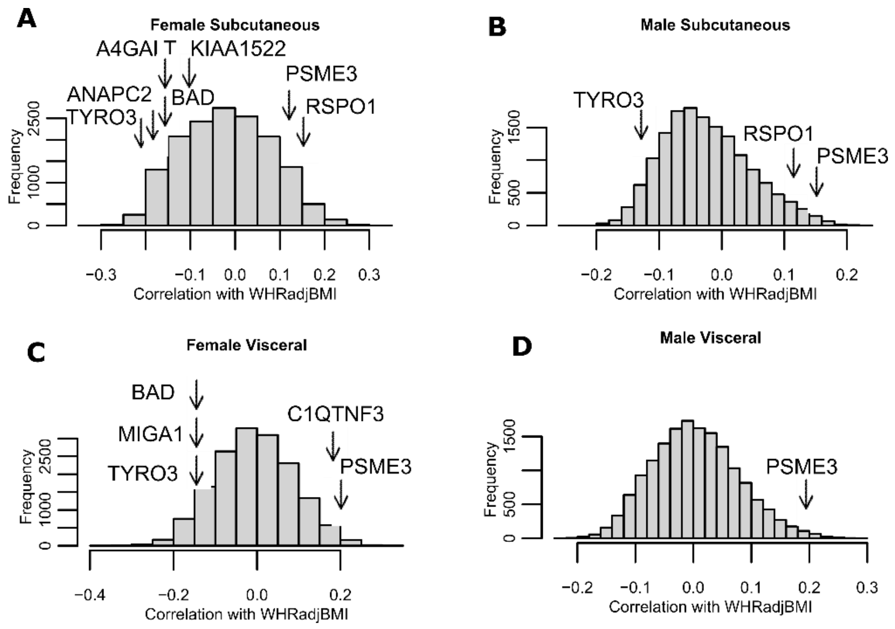


*Supplemental Figure 4: Gene expression correlations with WHRadjBMI in STARNET. Correlations with genes in (A) female subcutaneous (B) male subcutaneous, (C) female visceral, and (D) male visceral samples. Arrows indicate prioritized Wnt-related genes (Table 3) and mitochondrial-related genes (Table 4) whose correlation with WHRadjBMI is greater than 0.1.*

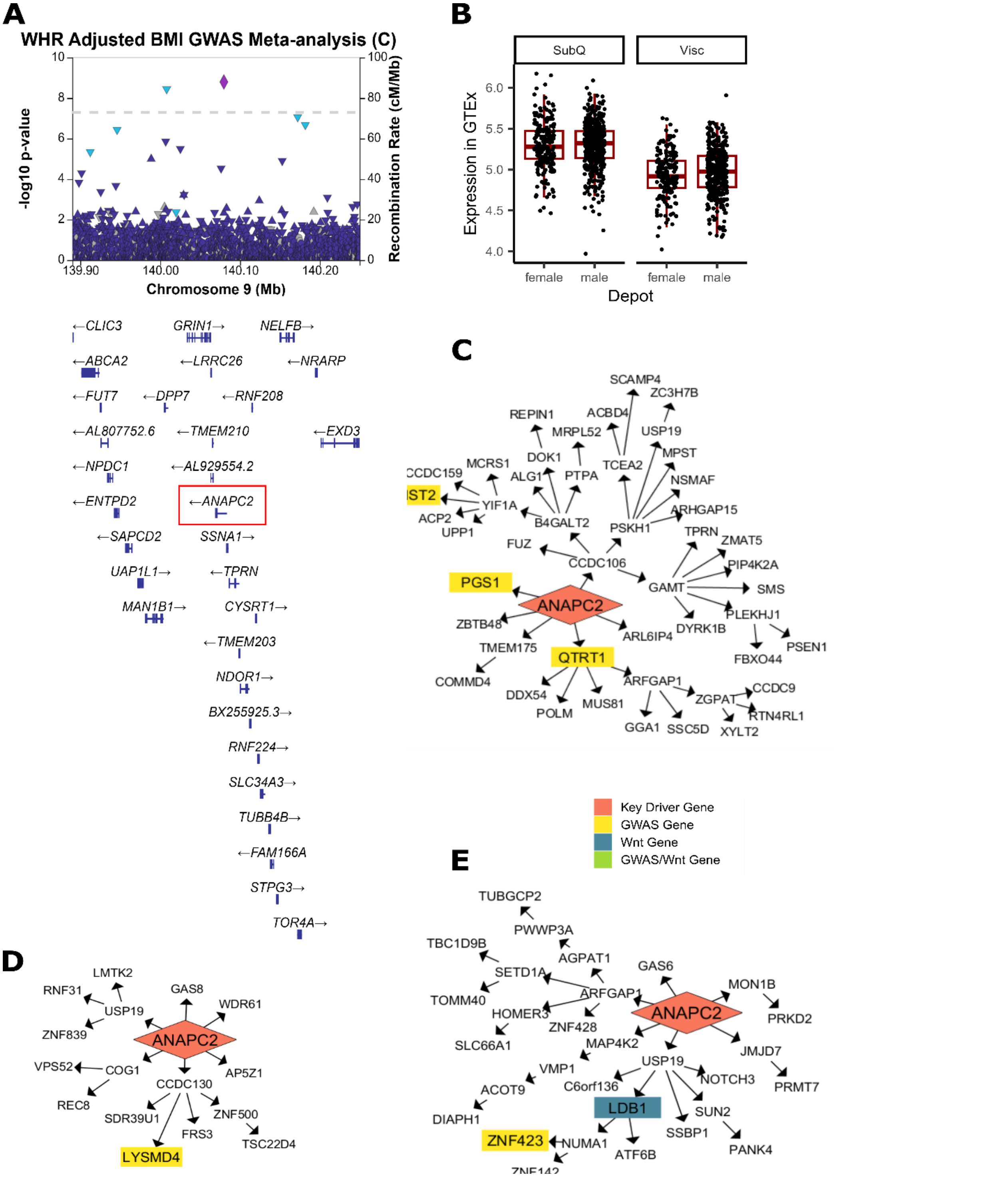


*Supplemental Figure 5- Additional evidence of ANAPC2 involvement in $WHR_{adjBMI}$ and Wnt signaling. (A) A significant GWAS signal is detected near ANAPC2 in $WHR_{adjBMI}$ GWAS meta-analysis[1]. (B) In GTEx, both males and females show higher expression of ANAPC2 in subcutaneous fat depots over visceral depots, though the change is non-significant after p-value adjustment. (C) ANAPC2 is a key driver that regulates downstream $WHR_{adjBMI}$ GWAS genes in the STARNET subcutaneous male network, (D) the GTEx*

*subcutaneous male network, and (E) regulates both $WHR_{adjBMI}$ GWAS genes and Wnt genes in STARNET female subcutaneous.*

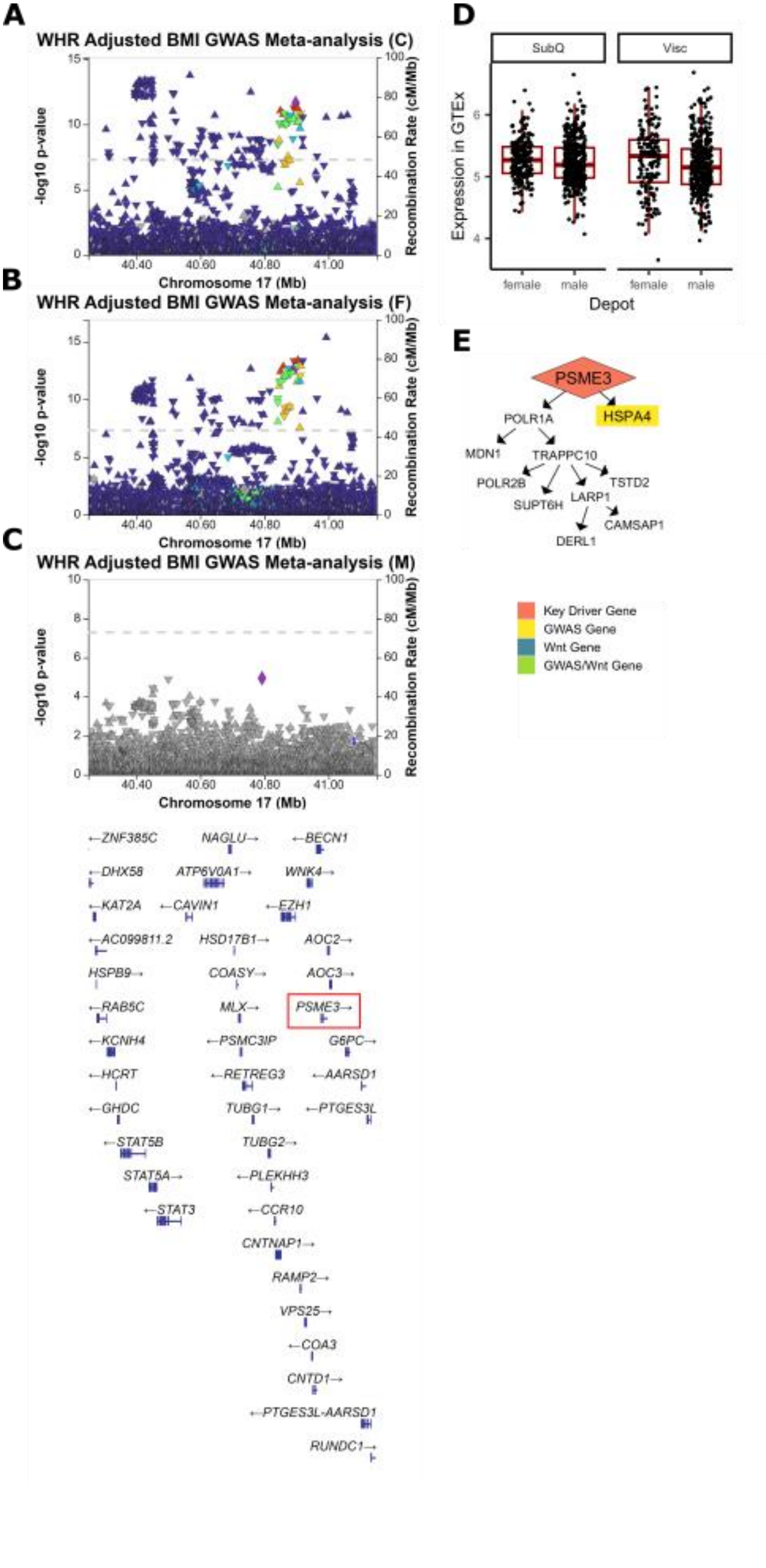
A
WHR Adjusted BMI GWAS Meta-analysis (C)
-log10 p-value
Recombination Rate (cM/Mb)
Chromosome 17 (Mb)
B
WHR Adjusted BMI GWAS Meta-analysis (F)
-log10 p-value
Recombination Rate (cM/Mb)
Chromosome 17 (Mb)
C
WHR Adjusted BMI GWAS Meta-analysis (M)
-log10 p-value
Recombination Rate (cM/Mb)
Chromosome 17 (Mb)
ZNF385C
NAGLU
BECN1
DHX58
ATP6V0A1
WNK4
KAT2A
CAVIN1
EZH1
AC099811.2
HSD17B1
AOC2
HSPB9
COASY
AOC3
RAB5C
MLX
PSME3
KCNH4
PSMC3IP
G6PC
HCRT
RETREG3
AARSD1
GHDC
TUBG1
PTGES3L
STAT5B
TUBG2
STAT5A
PLEKHH3
STAT3
CCR10
CNTNAP1
RAMP2
VPS25
COA3
CNTD1
PTGES3L-AARSD1
RUNDC1
D
SubQ
Visc
Expression in GTEx
female
male
female
male
Depot
E
PSME3
HSPA4
POLR1A
MDN1
TRAPPC10
POLR2B
TSTD2
SUPT6H
LARP1
CAMSAP1
DERL1
Key Driver Gene
GWAS Gene
Wnt Gene
GWAS/Wnt Gene

*Supplemental Figure 6- Additional evidence of PSME3 involvement in $WHR_{adjBMI}$. (A) A significant GWAS signal is detected near PSME3 in $WHR_{adjBMI}$ GWAS meta-analysis[1]. (B) This same signal is strongly associated with $WHR_{adjBMI}$ in the female-specific GWAS meta-analysis, (C) but is not present in the male-specific GWAS. (D) In GTEx, there are no significant changes in PSME3 gene expression between depots or sexes. (E) PSME3 regulates $WHR_{adjBMI}$ GWAS gene in the GTEx visceral female network.*

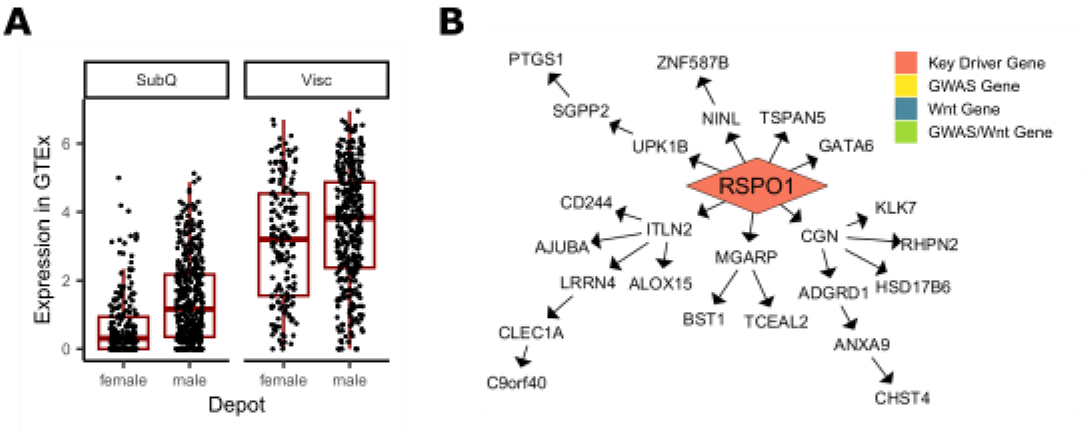


*Supplemental Figure 7- Additional evidence of RSPO1 involvement in $WHR_{adjBMI}$. (A) In GTEx, both sexes show higher expression of RSPO1 in visceral fat depots over subcutaneous depots (adjusted.P = 5.1e-21), and males show higher expression than females in both depots (adjusted.P = 4.9e-57). (B) RSPO1 is a key driver that regulates many downstream genes in the STARNET female visceral network.*

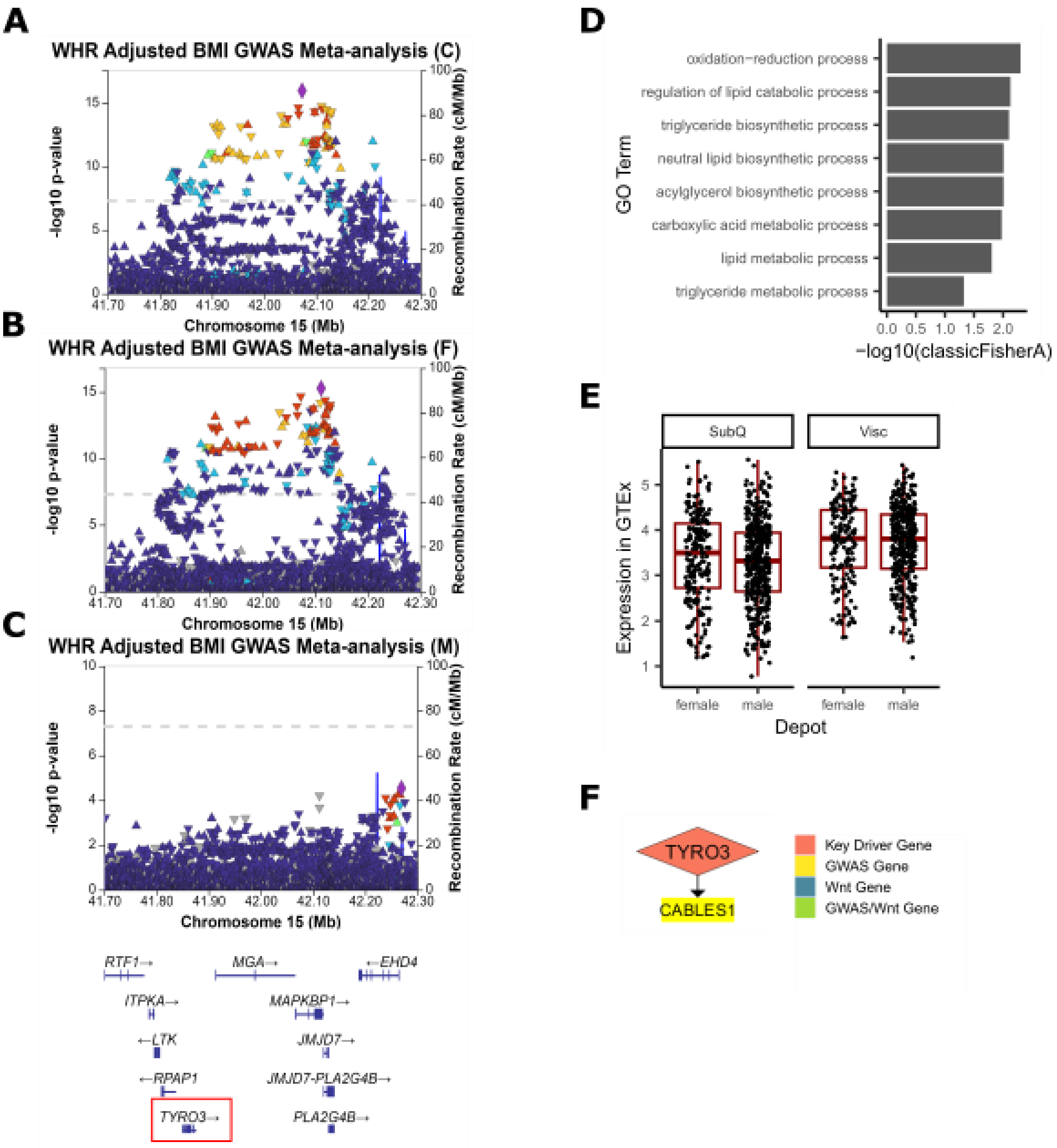


*Supplemental Figure 8- Additional evidence of TYRO3 involvement in $WHR_{adjBMI}$. (A) A significant GWAS signal is detected near TYRO3 in $WHR_{adjBMI}$ GWAS meta-analysis[1]. (B) This same signal is strongly associated with $WHR_{adjBMI}$ in the female-specific GWAS meta-analysis, (C) but is not present in the male-specific GWAS. (D) In GTEx subcutaneous male networks, the genes downstream of TYRO3 are significantly enriched for lipid-related biological processes. (E) In GTEx, both males and females show higher expression of TYRO3 in visceral fat depots over subcutaneous depots, though the change is non-significant after p-value adjustment. (F) TYRO3 regulates one $WHR_{adjBMI}$ GWAS gene in the STARNET subcutaneous male network.*

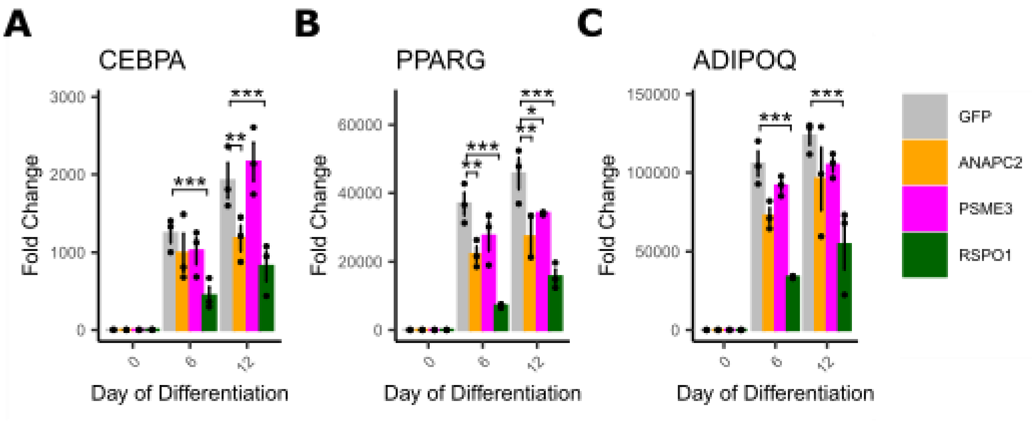


*Supplemental Figure 9- Markers of differentiation increase over time. Gene expression of (A) CEBPα, (B) PPARγ, (C) ADIPOQ increase over time, with some significant differences between cells overexpressing genes of interest. n = 3 replicates used in all assays. Differences between groups were determined using 1-way ANOVA within each timepoint by gene (Gene of Interest vs NTCcontrols). All post-hoc tests were performed using pooled t-test with Dunnett's adjustment. Adjusted p-values shown with * (*** = adj.P < 0.001,* = adj.P < 0.05, # = adj.P < 0.1).*

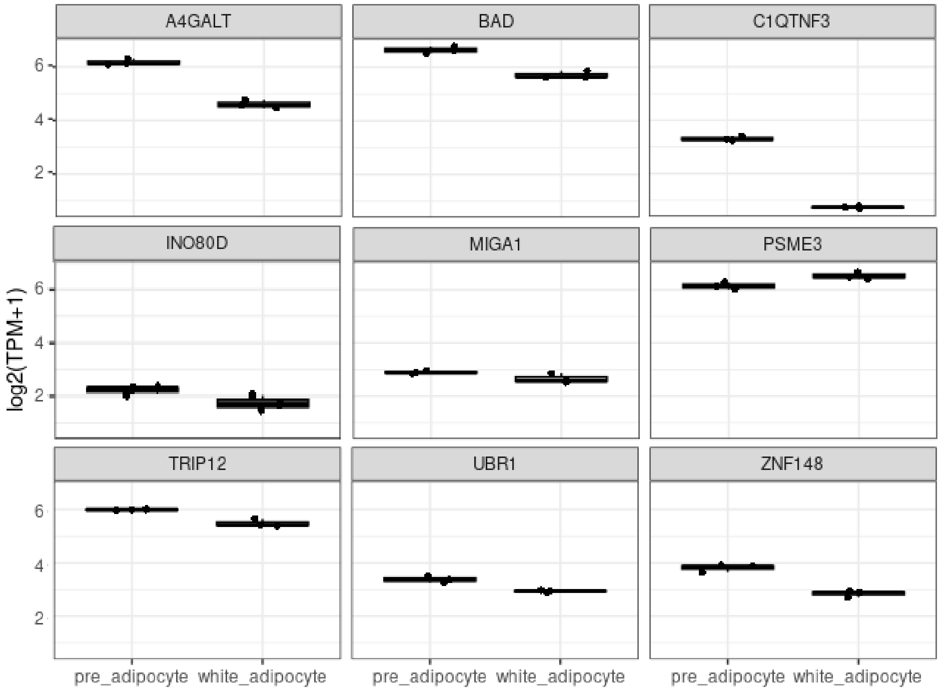


*Supplemental Figure 10- Mitochondrial candidate key driver expression in primary adipocyte cells.*

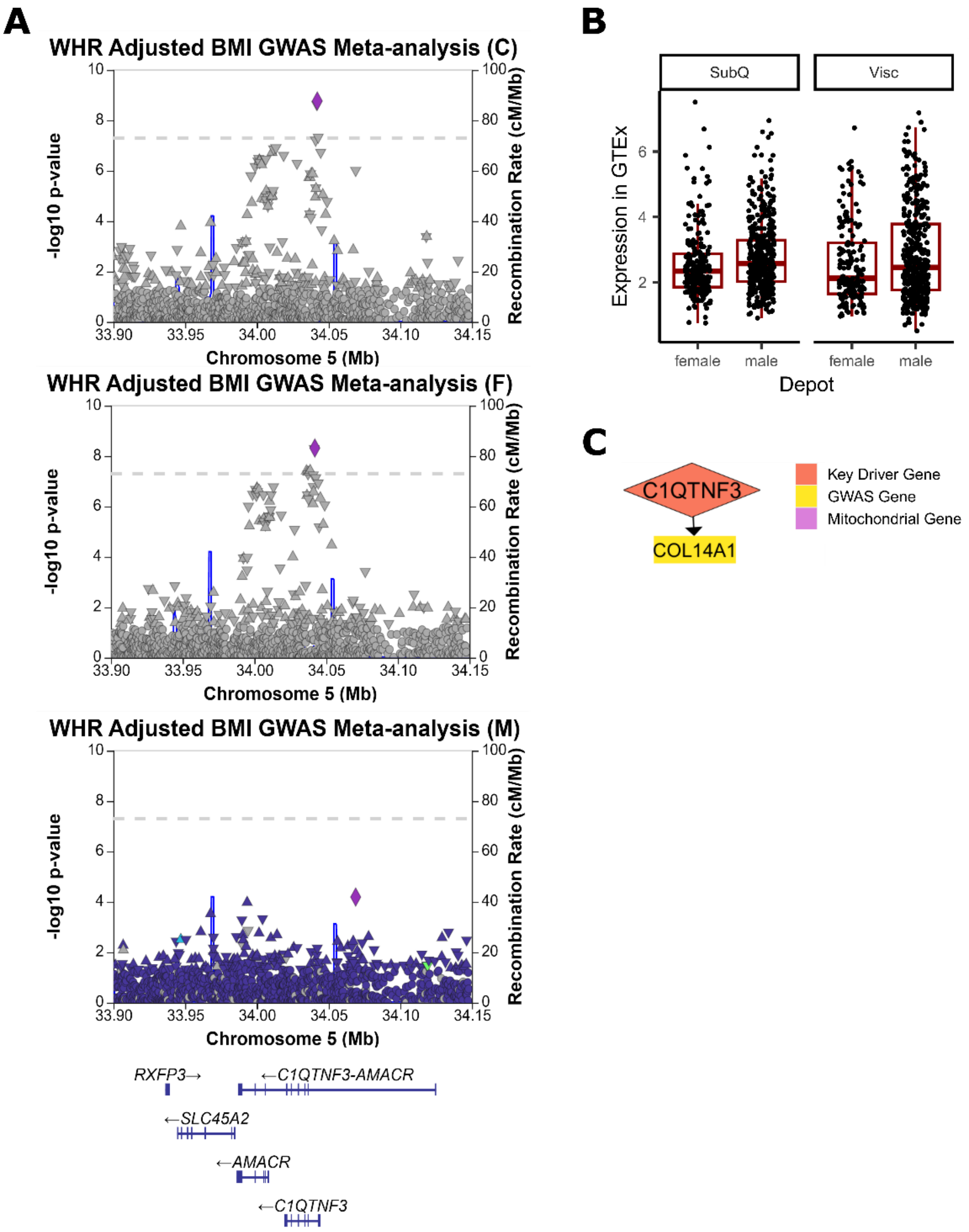


*Supplemental Figure 11- Additional evidence of C1QTNF3 involvement in $WHR_{adjBMI}$. (A) A significant GWAS signal is detected near C1QTNF3 in $WHR_{adjBMI}$ GWAS meta-analysis[1]. This same signal is strongly associated with $WHR_{adjBMI}$ in the female-specific GWAS meta-analysis, but is not present in the male-specific GWAS. (B) In GTEx, both males and females show higher expression of C1QTNF3 in visceral fat depots over subcutaneous depots, though the change is non-significant after p-value adjustment. (C) C1QTNF3 regulates one $WHR_{adjBMI}$ GWAS gene in the STARNET subcutaneous male network.*

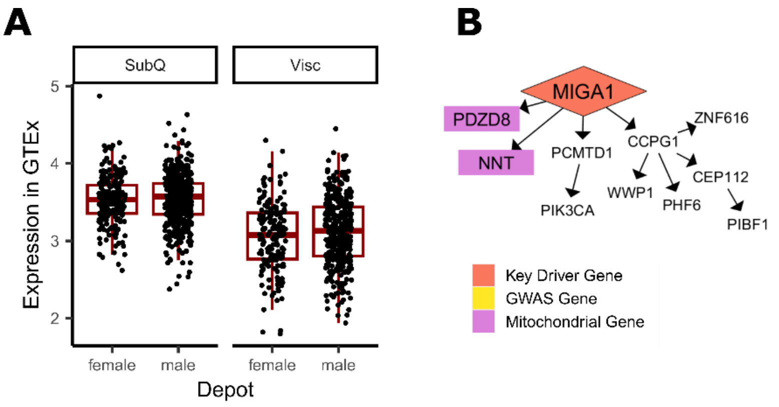


*Supplemental Figure 12- Additional evidence of MIGA1 involvement in $WHR_{adjBMI}$. (A) In GTEx, both males and females show higher expression of MIGA1 in visceral fat depots over subcutaneous depots, though the change is non-significant after p-value adjustment. (B) MIGA1 regulates mitochondrial genes in the STARNET subcutaneous male network.*

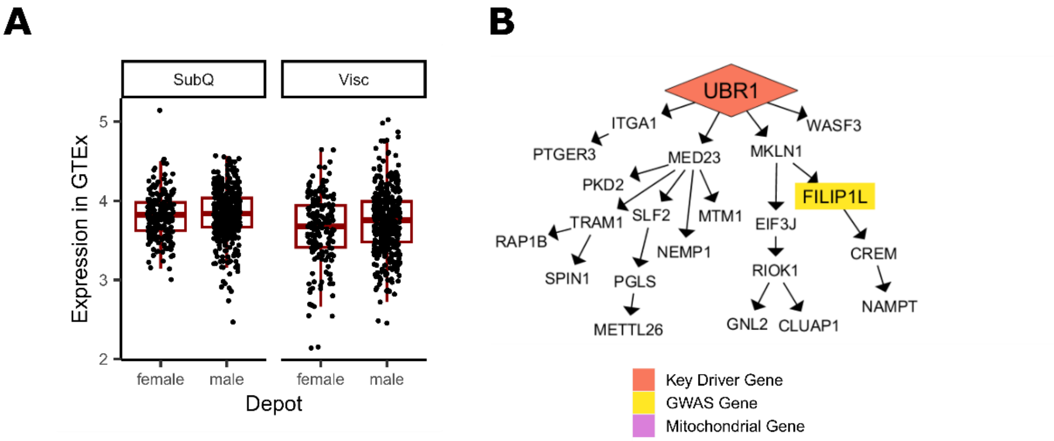


*Supplemental Figure 13- Additional evidence of UBR1 involvement in $WHR_{adjBMI}$. (A) In GTEx, both males and females show higher expression of UBR1 in visceral fat depots over subcutaneous depots, though the change is non-significant after p-value adjustment. (B) UBR1 regulates one $WHR_{adjBMI}$ GWAS genes in the STARNET subcutaneous male network.*

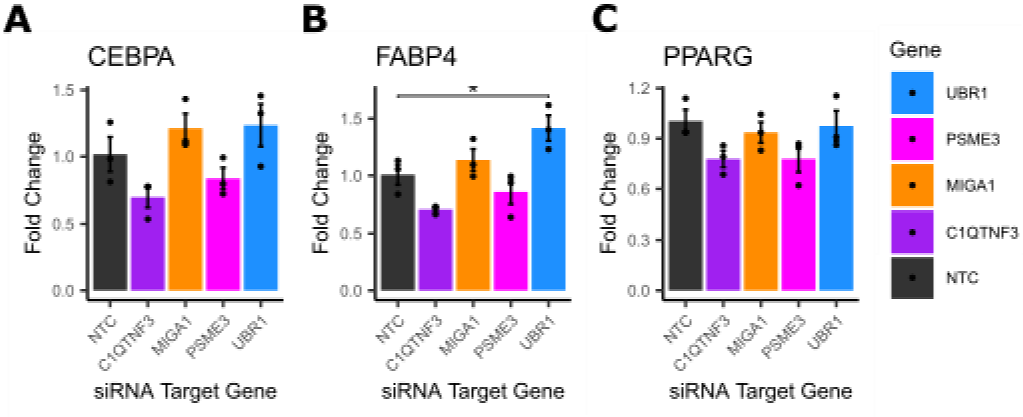


*Supplemental Figure 14- Markers of mature adipocytes FABP4 increases when mitochondrial key driver gene UBR1 is knocked down. Gene expression of (A) CEBPα, (B) PPARɣ, (C) FAPB4 in mature adipocytes treated with siRNA for the indicated gene. n = 3-6 replicates used in all assays. Differences between groups were determined using 1-way ANOVA by gene (Gene of Interest vs NTCcontrols). Post-hoc tests were performed using pooled t-test with Dunnett's adjustment. Adjusted p-values shown with * (*** = adj.P < 0.001,* = adj.P < 0.05, # = adj.P < 0.1).*

**Extended Bibliography 3: Evidence of Key Driver Involvement in Mitochondrial Activity**
